# Single-cell chromatin accessibility of developing murine pancreas identifies cell state-specific gene regulatory programs

**DOI:** 10.1101/2022.10.01.510484

**Authors:** Sean de la O, Zhe Liu, Sean Chang, Julie B. Sneddon

## Abstract

Numerous studies have characterized the existence of cell subtypes, along with their corresponding transcriptional profiles, within the developing mouse pancreas. The upstream mechanisms that initiate and maintain gene expression programs across cell states, however, remain largely unknown. Here, we generate single-nucleus ATAC-Sequencing data of developing murine pancreas and perform an integrated, multi-omic analysis of both chromatin accessibility and RNA expression to describe the chromatin landscape of both the developing epithelium and mesenchyme at E14.5 at single-cell resolution. We identify candidate transcription factors regulating cell fate and construct gene regulatory networks of active transcription factor binding to regulatory regions of downstream target genes. This work serves as a valuable resource for the field of pancreatic biology in general and contributes to our understanding of lineage plasticity among endocrine cell types. In addition, these data identify which epigenetic states should be represented in the differentiation of stem cells to the pancreatic beta cell fate in order to best recapitulate *in vitro* the gene regulatory networks that are critical for progression along the beta cell lineage *in vivo*.

## 1. Introduction

Development of the mammalian pancreas requires the coordination of multiple cell lineages over time, culminating in the generation of a highly branched, mature organ consisting of both an exocrine and endocrine compartment. Specification of the murine pancreas begins at embryonic day (E) 8.5 with the expression of the transcription factor (TF) pancreatic and duodenal homeobox 1 (*Pdx1*) in a focal region of the endoderm-derived primitive foregut [1, 2]. These Pdx1(+) cells give rise to all of the epithelial lineages of the pancreas (duct, endocrine, and acinar) [3] and by E9 evaginate into the surrounding mesenchyme and begin to form a stratified epithelium. As branching morphogenesis progresses, regionalization of the epithelium results in the formation of both trunk and tip domains by E12.5. Cells located at the tip, marked by expression of *Cpa1*, serve as multipotent progenitors that give rise to all three epithelial cell types until E13.5, at which point they undergo fate restriction to only give rise to acinar cells [3–5]. Epithelial cells located in the trunk give rise to either ductal or endocrine lineages, a fate choice dependent on levels of Notch signaling [5].

Endocrine progenitor (EP) cells derive from a subset of ductal epithelial cells that experience lower levels of Notch and then activate expression of the TF neurogenin3 (*Neurog3*) [5]. *Neurog3* expression marks early EP cells, which give rise to the main hormone-producing endocrine cells in the pancreas: alpha, beta, delta, and gamma [3, 6]. Gene knockout studies in mice revealed that the expression of a number of TFs that are critical for differentiation and maintenance of pancreatic endocrine lineages, such as paired box gene 4 (*Pax4*) and 6 (*Pax6*), neurogenic differentiation 1 (*Neurod1*), and LIM-homeodomain protein Islet 1 (*Isl1*), is dependent on *Neurog3* [6].

Endocrine cell identity is specified and maintained by a complex network of TFs, many of which play dynamic roles throughout developmental time [7]. For instance, early in development *Pdx1* is required for specification of pancreatic progenitors, but later in development it is also important for the generation of beta cells and for the maintenance of beta cell identity [8, 9]. Along with *Pdx1* and *Pax4*, the TFs NK2 homeobox 2 (*Nkx2-2*) and NK6 homeobox 1 (*Nkx6-1*) are critical factors for beta cell differentiation, while aristaless related homeobox (*Arx*) is essential for alpha cell differentiation. *Arx* and *Pax4* play mutually opposing roles in the differentiation of alpha and beta cells, with *Arx* promoting the generation of alpha at the expense of beta and delta cells [10] and *Pax4* regulating the decision towards beta and delta at the expense of alpha and epsilon cell fate [10–12]. When both *Arx* and *Pax4* are lost, delta cells persist but both alpha and beta cells are lost [11]. Expression of *Nkx2-2* and *Nkx6-1* follows that of *Pdx1* in early pancreatic progenitors, then becomes progressively restricted to endocrine cells [13, 14]. Deletion of *Nkx2-2* results in a significant reduction of the four major endocrine cell types and an increase in ghrelin-producing epsilon cells [12, 14]. *Nkx6-1* functions downstream of *Nkx2-2* and is necessary for beta cell neogenesis through the maintenance and/or expansion of beta cell precursors following *Neurog3* expression, but prior to the production of insulin, while later it is lost from developing alpha cells [13, 15].

Extrinsic signals derived from non-epithelial cells are also important in guiding pancreatic organogenesis. Early pioneering work using pancreatic explants *ex vivo* showed that when E11 epithelial buds were cultured without their surrounding mesenchymal tissue, epithelial growth and differentiation were arrested [16]. More recently, genetic ablation studies have demonstrated the requirement for pancreatic mesenchyme for expansion of the pool of early pancreatic progenitor cells early in development and for proliferation of differentiated cells later in development [17, 18]. Although the pancreatic mesenchyme is broadly appreciated as playing an important role in pancreatic organogenesis, however, it is still not well understood whether there exist biologically relevant sub-populations of mesenchyme with distinct lineages and/or functional roles.

Recent single-cell RNA-Sequencing (scRNA-Seq) studies have highlighted previously unappreciated levels of cellular heterogeneity among the epithelial cells of the developing murine pancreas, within the endocrine compartment in particular [19–23]. Although relatively less attention has been given to elucidating potential cellular heterogeneity within the mesenchymal compartment, evidence from scRNA-Seq and classical genetic lineage tracing experiments suggests that transcriptionally distinct mesenchymal cell types also exist during development [19,20,24]. As a result of this body of work, we now have a greater understanding of the transcriptomic cues governing cell states across pancreatic development, but we still lack an understanding of the upstream epigenetic features that regulate cell fate decisions. In particular, integration of gene expression data and chromatin accessibility data would permit identification of active transcription factor binding to accessible chromatin within a given cell type.

In recent years, Assay for Transposase-Accessible Chromatin followed by Sequencing (ATAC-Seq) has been developed to profile genome-wide chromatin accessibility for epigenetic analysis in a given cell type or tissue [25]. This technique has been applied to sorted populations of endocrine cells from the murine pancreas to investigate the chromatin landscape of developing EP cells [20, 26]. These studies, however, lacked single-cell resolution to capture the chromatin states of the various subpopulations of developing endocrine cells that have been described [19–23]. More recently, single-nucleus ATAC-Seq (snATAC-Seq) has emerged as a technology to provide insights into chromatin accessibility at single-cell resolution [27, 28]. snATAC-Seq has been used to profile the chromatin landscape of many developing tissue types and has revealed cell-type specific *cis*- and *trans*-regulatory elements governing gene expression and cell fate decisions [29–33]. Furthermore, integration of scRNA- and snATAC-Seq data for multi-omic analysis permits refinement of expressed TF to a further parsed subset that are likely binding TF motifs in open regions of chromatin and actively regulating expression of downstream target genes.

Here, we generate snATAC-Seq data of developing murine pancreas and perform an integrated multi-omic analysis of both chromatin accessibility and RNA expression. We describe at single-cell resolution the chromatin landscape of the developing epithelium at E14.5, a stage at which the dynamic processes of expansion, differentiation, and morphogenesis are actively underway. We identify candidate TFs regulating transitions across the endocrine lineages and construct gene regulatory networks (GRNs) of active TFs binding to regulatory regions of downstream target genes. Additionally, we generate an snATAC-Seq dataset of developing pancreatic mesenchyme, which to our knowledge represents the first ATAC-Seq dataset (bulk or single-nucleus) of this cell type. We believe these datasets and analyses will serve as a valuable resource for the field of pancreatic biology in general, and will contribute to our understanding of lineage plasticity among endocrine cell types. In addition, these data will serve as a reference as to which epigenetic states should be represented in the differentiation of stem cells to the pancreatic beta cell fate in order to best recapitulate *in vitro* the gene regulatory networks that are critical for progression along the beta cell lineage *in vivo*.

## 2. Methods

### 2.1. Animal studies

All mouse procedures were approved by the University of California, San Francisco (UCSF) Institutional Animal Care and Use Committee (IACUC). Mice were housed in a 12-hour light-dark cycle in a controlled temperature climate. Noon of the day of a vaginal plug was considered embryonic day (E)0.5. eFev-EYFP (ePet1-EYFP) mice were kindly donated by Dr. Evan Deneris, and have been previously described [34, 35]. Mice were maintained on a C57BL/6J background. Wildtype C57BL/6J mice used for breeding and for the whole pancreas sample were obtained from The Jackson Laboratory. Genotyping of eFev-EYFP mice was conducted on tail DNA, with forward primer TGCGATGGGAAGATAAGAGGGG and reverse primer GAAGTTCACCTTGATGCCGTTC.

### 2.2. Histology, immunofluorescence, and imaging

E14.5 pancreata were dissected in ice cold 1x PBS, then fixed in 4% paraformaldehyde (PFA) overnight at 4C. After washing three times in 1x PBS, tissues were preserved in 30% sucrose in PBS at 4C overnight and then embedded in Optimal Cutting Temperature (O.C.T.) compound (Tissue-Tek) and flash frozen prior to sectioning at 10 µm thickness.

For immunofluorescence staining, cryosections were washed 3 times in 1x PBS, permeabilized in 0.5% triton X-100 in PBS (PBT) for 10 minutes at room temperature (RT), and then blocked with 5% normal donkey serum (NDS) in 0.1% PBT for 1 hour. Sections were stained overnight at 4C using primary antibodies against GFP (1:500, Abcam Cat. ab13970) and Chga (1:250, Abcam Cat. ab15160). The next day, sections were washed three times in 1x PBS and then incubated with species-specific Alexa 488-, 555-, or 647-conjugated secondary antibodies and DAPI in 5% NDS in 0.1% PBT for 1 hour at RT. Sections were washed three times in 1x PBS and covered in Fluoromount-G mounting solution (SouthernBiotech, Cat. 0100-01).

Images were captured with an SP8 Leica confocal laser scanning microscope. Maximum intensity Z-projections were then prepared using Image J software [36].

### 2.3 *In situ* hybridization

Multiplexed *in situ* hybridization/immunofluorescence was performed with RNAscope technology using probes purchased from Advanced Cell Diagnostics, Inc. Probes against mouse *Fev* (Cat. 413241) and *Ngn3* (Cat. 422401) were used according to the manufacturer’s instructions for the RNAscope multiplex fluorescent detection V2 kit (Advanced Cell Diagnostics, Inc., Cat. 323110). 10 µm thick cryosections were brought to RT, washed with PBS to remove O.C.T., and treated with hydrogen peroxide and proteinase III. Tissue was hybridized with the probe mixture for 2 hours at 40C. Hybridization signals were amplified via sequential hybridization of amplifier AMP1, AMP2, and AMP3 and label probes Opal 570 (1:1500, PerkinElmer, Cat. FP1488001KT), Opal 650 (1:1500, PerkinElmer, Cat. FP1496001KT), and Opal 690 (1:1500, PerkinElmer, Cat. FP1497001KT).

Following signal amplification of the target probes, sections were incubated in 1x blocking buffer for 1 hour at RT, followed by staining with primary antibodies against GFP (1:500, Abcam Cat. ab13970). The next day, sections were washed three times with 1x PBS and then incubated with species-specific Alexa 488- or Alexa 555-secondary antibodies and DAPI in 5% NDS in 0.1% PBT for 1 hour at RT. Sections were then washed three times in 1x PBS, mounted with ProLong Gold Antifade Mountant (Invitrogen, Cat. P36930), and stored at 4C prior to imaging. Optical sectioning images were taken with a Leica confocal laser scanning SP8 microscope equipped with white light sources. 10 steps X 1 mm thickness Z-sections were captured for each imaging area.

### 2.4. Dissociation and sorting of murine pancreas tissue for quantitative RT-PCR

E14.5 pancreata were dissected from embryos of pregnant eFev-EYFP dams, and kept in separate wells of a 96-well plate. EYFP fluorescence was assessed under a microscope to confirm the genotype of each pancreas. Pancreata with EYFP fluorescence (EYFP(+)) were then pooled together and transferred to a 1.5 ml microcentrifuge tube, then dissociated into single cells by incubating with 250 ul of TrypLE Express dissociation reagent (Gibco, Cat. 12604013)) at 37C for 20 minutes, with pipet trituration at 5 minute intervals. Dissociation was neutralized with FACS buffer (10% FBS + 2 mM EDTA in phenol-red free HBSS), and the single-cell suspensions were passed through 30 µm cell strainers.

Cells were stained with SYTOX Blue dead cell stain (Invitrogen, Cat. S34857) to remove dead cells, then with a PE-conjugated antibody against mesenchymal marker CD140a (1:50; eBioscience Cat. 12-1401-81) and an APC-conjugated antibody against epithelial marker CD326/EpCam (1:50; eBioscience Cat. 17-5791-82) at 4C for 30 minutes. Stained cells were washed twice in FACS buffer and sorted using a BD FACSAria II cell sorter (BD Biosciences). After size selection to remove debris and doublets and sorting on SYTOX Blue negative (live) events, cells were further subgated on CD140a(-)/CD326(+) (epithelial) cells and then on EYFP fluorescence.

RNA was extracted from EYFP(-), EYFP-low, and EYFP-high sorted cells with the RNeasy Mini Kit (Qiagen, Cat. 74106). Reverse transcription was performed with the PrimeScript High Fidelity RT-PCR Kit (Takara, Cat. R022A). RT-PCR was run on a 7900HT Fast RT-PCR instrument (Applied Biosystems) with Taqman probes for *Fev* (assay ID: Mm00462220_m1, Cat. 4331182) and *GAPDH* (assay ID: Mm99999915_g1, Cat. 4331182) in triplicate. Data were normalized to *GAPDH*. Error bars represent standard error of the mean (SEM).

### 2.5. Dissociation and sorting of murine pancreas tissue for snATAC-Seq

For the whole pancreas sample, E14.5 C57BL/6J embryonic pancreata (n=10) were dissected from 3 litters and pooled into a single 1.5 ml microcentrifuge tube. For the eFev-EYFP sample, E14.5 pancreata (n=15) were dissected from embryos of two pregnant eFev-EYFP dams, and kept in separate wells of a 96-well plate. EYFP fluorescence was assessed under a microscope to confirm the genotype of each pancreas. Pancreata (n=5) with EYFP fluorescence (EYFP(+)) were then pooled together, pancreata (n=10) without EYFP fluorescence (EYFP(-)) were pooled together (as a negative control), and each sample was transferred to a separate 1.5 ml microcentrifuge tube.

The whole pancreas, EYFP(+), and EYFP(-) samples were dissociated into single cells by incubating with 250 ul per sample of TrypLE Express dissociation reagent (Gibco, Cat. 12604013)) at 37C for 20 minutes, with pipet trituration at 5 minute intervals. Dissociation was neutralized with FACS buffer (10% FBS + 2 mM EDTA in phenol-red free HBSS) and the single-cell suspensions were passed through 30 µm cell strainers.

All samples were stained with SYTOX Blue dead cell stain (Invitrogen, Cat. S34857) to remove dead cells. Cells were washed twice in FACS buffer and sorted using a BD FACSAria II cell sorter (BD Biosciences). After size selection to remove debris and doublets, all cells were sorted on SYTOX Blue negative (live) events, and the EYFP(+) and EYFP(-) samples were further subgated on EYFP fluorescence. Live cells from whole pancreas and live EYFP(+) cells from the EYFP(+) sample were collected into separate tubes containing 1x FACS buffer and immediately subjected to extraction of nuclei as described below.

### 2.6. Extraction of nuclei

All buffers (e.g., 0.1x lysis buffer, lysis dilution buffer, and wash buffer) were freshly prepared according to the 10x Genomics Demonstrated protocol (CG000212 RevC), and maintained at 4C. Nuclei were isolated from 25,000 cells from whole pancreas or from Fev-high (EYFP(+)) cells using the demonstrated protocol. In brief, sorted cells were added to a 2 ml microcentrifuge tube and centrifuged at 500 rcf for 5 minutes at 4C. All supernatant was removed without disrupting the cell pellet. 100 ul chilled 0.1x lysis buffer was then added and pipetted 5 times to fully mix the buffer with the cells, then incubated for 3 minutes on ice to achieve full cell lysis. 1 ml chilled wash buffer was added to the lysed cells to terminate the lysis. Lysed cells were centrifuged at 500 rcf for 5 minutes at 4C, and supernatant was gently removed. Nuclei were resuspended in 50 ul wash buffer, transferred to a 200 ul tube, and spun down and resuspended in 10 ul 1x Nuclei buffer (10x Genomics, Part Number 2000153). 2 ul of the suspension was loaded onto a hemacytometer to determine the concentration of nuclei and simultaneously assess nucleus quality. 12,500 high-quality nuclei from the whole pancreas sample and 5,000 high-quality nuclei from the eFev-EYFP sample were then used for downstream library construction and sequencing.

### 2.7. snATAC-seq capture, library construction, and sequencing

Input nuclei were subjected to transposition, partitioning, and library construction using 10x Genomics Chromium Next GEM Single Cell ATAC Reagent Kit v1.1 Chemistry, according to the manufacturer’s instructions. An Agilent Fragment Analyzer was used for assessing the fragment distribution of both the whole pancreas and eFev-EYFP libraries, which were run on the Illumina NovaSeq 6000 platform.

### 2.8 Clustering of murine scRNA-Seq data

For clustering of murine scRNA-Seq data for integration with our snATAC-Seq data, we applied the clustering algorithm CellFindR [37] to our previously published dataset of developing murine pancreas tissue [19]. 10x Genomics outputs of E14.5 pancreata were downloaded from the Gene Expression Omnibus (GEO) (GSE101099; samples GSM3140916, GSM3140919 and GSM3140920), and analyzed with Seurat v3.2.3. Seurat objects were created from each 10x output with Read10x() and CreateSeuratObject() and filtered to retain high quality cells (nFeature_RNA > 1250 and percent.mt < 7 for GSM3140916; nFeature_RNA > 1500 and percent.mt < 5 for GSM3140919 and GSM3140920). The datasets were then normalized and variable features calculated with NormalizeData() and FindVariableFeatures(), respectively. The samples were then integrated using Seurat’s standard batch correction method [38] () with SelectIntegrationFeatures(), FindIntegrationAnchors() and IntegrateData(). The integrated object was then scaled with ScaleData() and principal component analysis (PCA) performed with RunPCA(). UMAP dimensional reduction was calculated with RunUMAP() with dims = 1:30. Neighbors were found in the dataset with FindNeighbors() with dims = 1:30 and clustering performed with FindClusters(), resolution = 0.2. Next, broad cell types were manually annotated based on expression of known marker genes (i.e. *Col3a1* for mesenchyme) and used for subsequent iterative sub-clustering.

For iterative subclustering with CellFindR, each broad cell type (Mesenchyme, Mesothelium, Exocrine, and Endocrine) was subsetted individually. PCA, Neighbors and UMAP were recalculated as described above (Mesenchyme: dims 1:15; Mesothelium: dims 1:15; Exocrine: dims 1:10; and Endocrine: dims 1:10) and the first clustering resolution calculated with find_res() from CellFindR. Iterative subclustering was then performed with sub_clustering(). Subclusters that displayed characteristics of doublets (expressing markers of more than one broad group e.g., *Col3a1+*/*Cpa1+* acinar cells) or low quality (e.g. clustering based on high mitochondrial gene content) were manually removed.

### 2.9. snATAC-Seq analysis

FASTQ files were generated from raw sequencing reads using the bcl2fastq function from Illumina. BAM files and single-cell accessibility counts were generated using the cellranger-atac count function from Cell Ranger software, version 1.0.1. Reference genome used was Mus musculus assembly mm10, annotation gencode.vM17.basic. Files processed with Cell Ranger ATAC were then analyzed using ArchR (version 1.0.1) [39].

First, ArchR Arrow files were created with the ArrowFiles() function with default settings. An ArchR project was then created using both whole pancreas and EYFP(+) sorted cells with ArchRProject(). The project was then filtered for high quality nuclei (TSS enrichment >= 10 and number of fragments >= 3,000) and doublets removed with addDoubletScores() and filterDoublets(), resulting in a final dataset consisting of 15,003 total cells with a median number of 14,630 fragments per cell and a median TSS enrichment score of 14.673. Next, iterative LSI was performed with the addIterativeLSI() function, with clustering parameters of resolution = 0.2, sampleCells = 10,000, n.start = 10, varFeatures = 25000 and dimsToUse = 1:20. Clustering was performed with addClusters() with resolution = 0.1 and method = “Seurat”. UMAP dimensional reduction was performed with addUMAP(), with minDist = 0.5. Clusters were manually annotated based on the Gene Score of known marker genes with addImputeWeights() and then visualized by UMAP.

For epithelial analysis, epithelial (exocrine and endocrine) nuclei were subsetted based on accessibility of known marker genes (*Cpa1, Spp1, Chga*). Iterative LSI was recalculated with iterations = 2, resolution = 0.2, sampleCells = 5,000, n.start = 10, varFeatures = 25000 and dimsToUse = 1:15. Clustering was performed with resolution = 0.9, and UMAP recalculated with minDist = 0.5 and dimsToUse = 1:15. Clustered epithelial cells from the scRNA-Seq data described above were used for unconstrained integration with addGeneIntegrationMatrix(). Chromatin accessibility peaks were then called with Macs2 via ArchR with addGroupCoverages(), addReproduciblePeakSet() and addPeakMatrix(). Marker peaks within the epithelial compartment were calculated with getMarkerFeatures() using the “PeakMatrix”. For motif analysis within marker peaks, motif annotations were added with addMotifAnnotations() with the “cisbp” motif set and then calculated with peakAnnoEnrichment(). ChromVAR [40] analysis was performed with addBgdPeaks() and addDeviationsMatrix(). Correlated transcription factors were correlated between the “GeneIntegrationMatrix” (RNA expression from the unconstrained integration) and “MotifMatrix” (ChromVAR motif deviations) with correlateMatrices(), keeping TFs with a correlation > 0.5, padj < 0.01 and max delta greater than 0.5 of the upper quartile.

For pseudotime lineage calculations, we manually imputed the cell states for each cell lineage (Alpha and Beta) and computed the pseudotime values with addTrajectory().

For mesenchymal and mesothelial analysis, nuclei were subsetted based on accessibility of known marker genes (*Col3a1, Wt1*). Iterative LSI was recalculated with iterations = 2, resolution = 0.5, sampleCells = 2,500, n.start = 10, varFeatures = 25000 and dimsToUse = 1:15. Clustering was performed with resolution = 0.3, and UMAP recalculated with minDist = 0.3 and dimsToUse = 1:15. Clustered mesenchymal and mesothelial cells from the scRNA-Seq data described above were used for unconstrained integration. Peak calling, marker peak identification, motif analysis, ChromVAR analysis and transcription factor correlation were performed as described above.

### 2.10. Gene regulatory network analysis

Gene regulatory networks (GRNs) were constructed as described by Lyu et al., 2021 [31] (https://github.com/Pinlyu3/IReNA-v2). Candidate cluster-enriched genes were calculated with the scRNA-Seq dataset of epithelial or mesenchymal cells with Seurat’s FindAllMarkers() with min.pct = 0.1 and logfc.threshold = 0.25. DEGs were retained with an average logFC > 0 and padj < 0.01. DEGs were then mapped to specific cell types with the IReNA v2 function Process_DEGs_to_Celltypes().

Peak-to-gene linkage was performed with ArchRs addPeak2GeneLinks() function using dims = 1:15 for both epithelium and mesenchyme using the “GeneIntegrationMatrix” (integrated RNA Seq counts). Peak-to-gene links were then extracted with the IReNA v2 function Get_p2g_fun().

To identify potential *cis*-regulatory elements for each candidate gene, called correlated accessible regions (CARs), we separated the peak-to-gene links into three categories: TSS (when the peak lies within the transcription start site (TSS) for the gene), gene body (when the distance between peak and TSS is less than 100kb, and the peak-to-gene score calculated above is significant), or distal (when the peak is 100kb upstream or downstream of the TSS of correlated gene, and the peak-to-gene score is significant). These peak-to-gene links were then filtered to only include genes in the DEG list calculated above with Selection_peaks_to_one().

Next, we predicted the cell-type specific transcription factors binding in these CARs. We first took the snATAC fragments for each dataset (whole pancreas and Fev-high) and then extracted the fragments for each cell type. We converted these fragment lists to .BAM files and corrected the Tn5 insertion bias with TOBIAS [41] ATACorrect with default parameters except -- read_shift 0 0. We then converted the TOBIAS output bigwig files to GRanges with the IReNA v2 function Check_normalized_Signal(). Next, we calculated TF binding motifs in our peaks with motifmatcher (https://github.com/GreenleafLab/motifmatchr), filtering calculated TFs out from the motif analysis if they were not enriched in each cell type by the DEG analysis. Next, we calculated the NC (average bias-corrected Tn5 signal in the center of the motif), NL and NR (average bias-corrected Tn5 signal in the left and right flanking regions of the motif) scores with Calculate_footprint_celltypes() and filtered TFs with a score of NC < −0.1 and NL > 0.1 and NR > 0.1.

Next, we used MAGIC (Mining Algorithm for GenetIc Controllers) [42] to compute correlation between TF and target gene gene expression. We retained the top and bottom 2.5% of correlations for our downstream analysis.

Lastly, we constructed the cell-type specific GRNs. We combined the peak-target links from our third step with the cell-type specific TF-peak links from our fourth step with Reg_one_cells_RPC_MG(). We then classified these interactions as either activating or repressing with our TF-target gene interactions calculated above with Add_Cor_to_GRN_network_and_Filter(). We then identified feedback TF-TF pairs in our constructed GRN with FoundFeedBackPairs_new() and Process_the_Feedback_res().

## 3. Results

### 3.1. Single-nucleus ATAC-Sequencing of the developing murine pancreas

To investigate chromatin accessibility in the developing pancreas, we aimed to capture a broad range of cell types, including both epithelial and non-epithelial populations. In addition, we were specifically interested in profiling endocrine progenitor (EP) cells, but given their rare numbers we searched for a method to enrich for this population. We utilized ePet1-EYFP mice (referred to hereafter as eFev-EYFP, as the gene *Pet1* is also known as *Fev*), where EYFP expression is driven by a *Fev* enhancer [34, 35]. In previous work, we had identified *Fev* as a marker of an intermediate murine EP population downstream of the better-characterized Ngn3(+) population and upstream of differentiated, hormone-expressing endocrine cells [19].

As lineage reconstruction of scRNA-Seq data had revealed that this Fev-expressing EP population is likely the state at which endocrine lineage allocation occurs, we chose to enrich Fev(+) cells. Although previous work with this eFev-EYFP mouse line had validated that EYFP expression faithfully reflected *Fev* expression in brain tissue, similar confirmation had not yet been performed in the pancreas [34]. We performed dual *in situ* hybridization (ISH)/immunofluorescence (IF) staining of E14.5 eFev-EYFP pancreas tissue to evaluate the architecture of EYFP expression with respect to the expression of *Ngn3* and *Fev* transcripts, as well as Chromogranin A (Chga) protein, a marker of differentiated hormone-producing endocrine cells. Expression of *Ngn3* and *Fev* transcripts was mutually exclusive (**Figure 1A**), as expected from our previous work demonstrating by genetic lineage tracing and scRNA-Seq that Fev-expressing cells are downstream of an Ngn3(+) state [19]. In contrast, a significant fraction, but not all, of EYFP(+) cells were actively expressing *Fev* transcript (**Figure 1A**). In addition, we observed EYFP(+) cells also expressing Chga (**Figure S1A**). As expected, EYFP expression was only found in epithelial (E-cadherin(+)) cells, and mostly localized to ductal-like structures (**Figure S1A**). These data are consistent with a model of *Fev* expression in pancreatic EP cells in eFev-EYFP mice in which *Fev* transcript first begins to be expressed as *Ngn3* expression wanes, then expression of EYFP (under the control of the *Fev* enhancer) follows (**Figure 1B**). Persistence of EYFP in cells that no longer express *Fev* transcript likely reflects longer perdurance of EYFP fluorescent protein compared to *Fev* mRNA in these cells, similar to what has been observed for Ngn3-tdTomato [43], Ngn3-YFP [44]), and Ngn3-EGFP transgenic mice [45].

**Figure 1:**
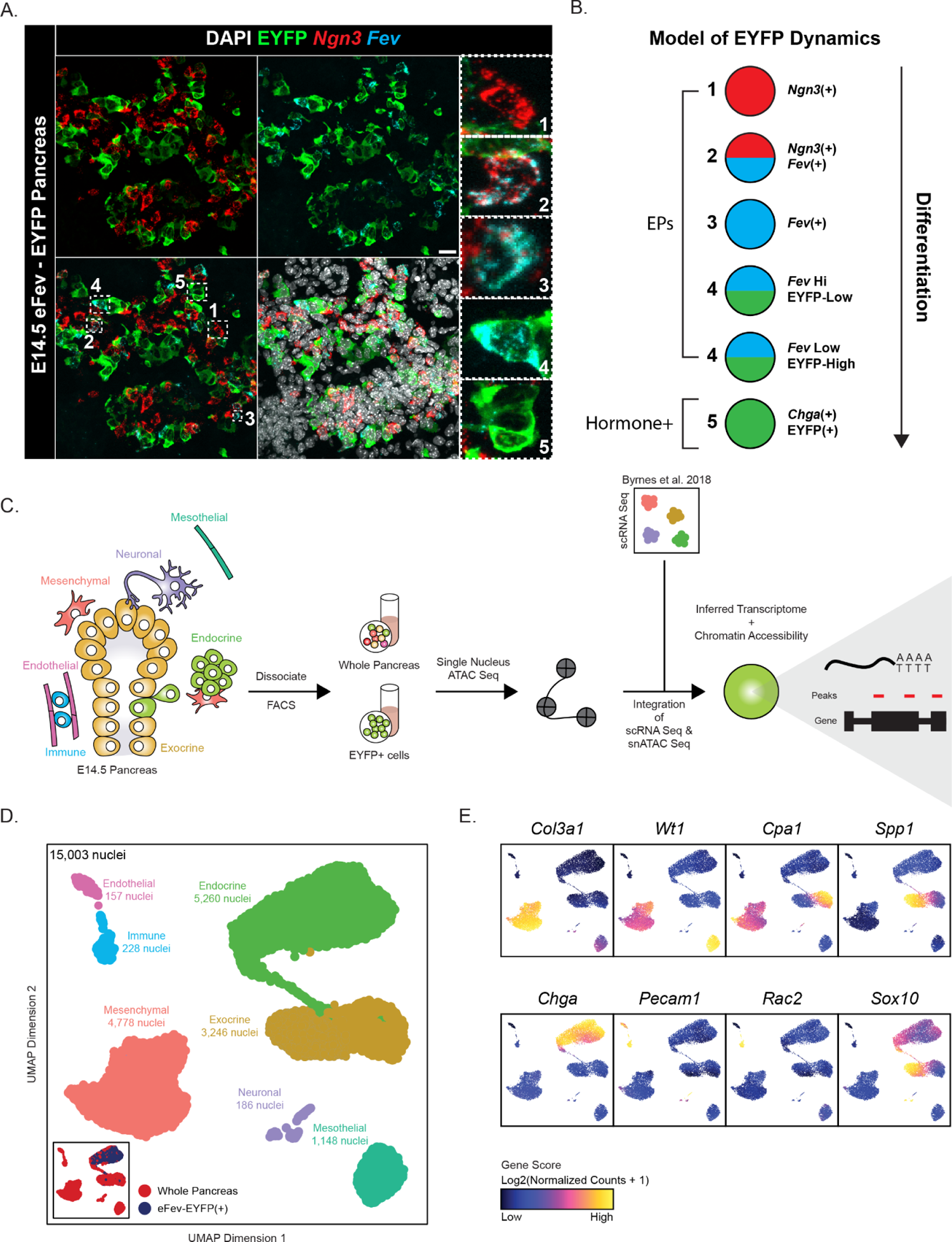
Single-nucleus ATAC-Seq of developing murine pancreas. (**A**) Multiplexed immunofluorescence and *in situ* hybridization staining of E14.5 pancreas from eFev-EYFP transgenic mouse embryos. *Neurog3* transcript is shown in red and marks early endocrine progenitors (EPs), *Fev* transcript is shown in cyan and marks intermediate EPs, and EYFP protein is shown in green. DAPI marks nuclei in white. Five selected regions of interest (ROI) are outlined by dashed white rectangles on the merged image and shown at higher magnification to the right. These ROIs highlight examples of cells that are 1) Ngn3+/Fev-/EYFP-; 2) Ngn3+/Fev+/EYFP-; 3) Ngn3-/Fev+/EYFP-; 4) Ngn3-/Fev+/EYFP+; and 5) Ngn3-/Fev-/EYFP+. Scale bar is 20uM. (**B**) Model of *Fev* and EYFP expression in EPs undergoing differentiation to a hormone-producing, Chromogranin A (Chga)-expressing state in eFev-EYFP reporter mice. Each circle represents a cell state across endocrine differentiation. (**C**) Overview of experimental approach for generating single-nucleus ATAC-Seq (snATAC-Seq) data. To enrich for Fev-high EPs, E14.5 eFev-EYFP murine pancreas was dissociated, and the resulting single cell suspension was subjected to FACS to enrich for EYFP+ epithelial cells (“EYFP+ Cells”). In parallel, E14.5 pancreata from control (C57BL/6J) embryos were dissociated and subjected to FACS to isolate all live cells (“Whole Pancreas”) to profile a broad spectrum of cell types, including non-epithelial cells. After subjecting samples to snATAC-Seq, data were then integrated with previously-published single-cell RNA-Sequencing (scRNA-Seq) datasets of E14.5 murine pancreas previously published by our laboratory [19]. (**D**) Uniform Manifold Approximation and Projection (UMAP) visualization of merged snATAC-Seq datasets from both Whole Pancreas and EYFP+ samples, comprising a total of 15,003 nuclei. Each dot represents a single cell, and each cell is colored according to cell type. Contribution of each sample (Whole Pancreas and eFev-EYFP+) to the total dataset is depicted in the inset, with the eFev-EYFP+ sample contributing only to the endocrine cluster as expected. (**E**) Feature plots depicting the Gene Scores (accessibility of the gene promoter plus the gene body) for some of the marker genes used to annotate the cell types in panel **D**.

We further validated the eFev-EYFP mouse line using fluorescence-activated cell sorting (FACS) and quantitative real-time polymerase chain reaction (qRT-PCR). Consistent with our IF staining, we observed little to no EYFP signal in cells that were negative for the epithelial marker EpCAM (**Figure S1B**). Within the population of cells positive for EpCam and negative for the mesenchymal marker CD140a, a bimodal distribution of EYFP signal was detected (**Figure S1B**). TaqMan qRT-PCR analysis revealed that the EYFP-low population had higher expression of *Fev* mRNA compared to EYFP-high cells (**Figure S1C**). The EYFP-low population thus likely corresponds to a stage in which EYFP expression is on the rise and *Fev* expression is still present, whereas the EYFP-high population likely represents a stage where EYFP has reached higher expression but *Fev* itself has begun to wane (**Figure 1B**). Thus, we selected this EYFP-low population, enriched for Fev(+) cells, for snATAC-Seq using the 10x Genomics platform (**Figure 1C, Figure S1D**). We additionally included a second sample in our analysis consisting of whole pancreas, to capture a broad range of cell types (**Figure 1C**).

Single cells were lysed to isolate nuclei, and chromatin was then subjected to the 10x Genomics pipeline and sequenced. The resulting dataset was analyzed with the computational package ArchR [39]. First, the datasets were filtered to retain high-quality nuclei by thresholding on the number of unique nuclear fragments, as well as the transcription start site (TSS) enrichment score (see Methods). This step enriches for cells displaying a high fraction of fragments that map to the TSS, versus other locations in the genome. Next, the datasets were subjected to doublet discrimination, resulting in a final dataset consisting of a combined total of 15,003 high-quality nuclei across the two samples. The data were then dimensionally reduced, clustered, and visualized in a 2D Uniform Manifold Approximation and Projection (UMAP) embedding (**Figure 1D**). As expected, cells from the eFev-EYFP(+) sample clustered only with endocrine cells from the Whole Pancreas sample (**Figure 1D**, inset), reflecting successful enrichment of endocrine cells from the eFev-EYFP mouse line and effective integration of the two datasets. Each cluster was annotated as corresponding to a specific cell type found within the developing pancreas based on the gene score (accessibility of the gene promoter plus the gene body) of the following marker genes: *Col3a1* to mark mesenchymal cells, *Wt1* for mesothelial cells, *Cpa1* and *Spp1 for* exocrine cells, *Chga* for endocrine cells, *Pecam1* for endothelial cells, *Rac2* for immune cells, and *Sox10* for neuronal cells (**Figure 1E, Supplemental table 1**).

### 3.2. Integration of single-cell transcriptional and chromatin accessibility data identifies epithelial heterogeneity in the developing murine pancreas

To reliably identify the heterogeneity of chromatin states within the epithelial cell types of the developing pancreas, we performed unconstrained integration of our snATAC-Seq data from all epithelial cells with E14.5 scRNA-Seq data previously published by our laboratory [19]. First, we computationally isolated the epithelial cells from the scRNA-Seq dataset (13,093 epithelial cells total) and performed iterative sub-clustering with the computational package CellFindR [37] to identify biologically relevant cell types. Next, we correlated the gene expression profiles of each of the cells within this scRNA-Seq dataset with the gene scores of each of the cells within our snATAC-Seq dataset. After identifying correlated cell pairs between the two datasets, cells in the snATAC-Seq dataset were assigned the cell type label, as well as the gene expression profile, of the cognate cell from the scRNA-Seq data.

This integration resulted in a final epithelial snATAC-Seq dataset comprised of 8,506 nuclei representing 10 distinct cell types, including Acinar, Ductal, Spp1(+)/Neurog3(+) double positive EPs, Neurog3(+) single positive EPs, Fev(+)/Chgb(+) intermediate progenitors, and Pdx1(+)/Mafb(+) beta cell precursors, as well as Alpha, Beta, Delta and Epsilon cells (**Figure 2A**). As expected, the sorted EYFP(+) cells contributed highly to the endocrine but not the acinar or ductal compartments of the overall dataset (**Figure 2A, inset**). The relative proportions of these annotated cell types in the snATAC-Seq dataset roughly matched the proportions of the epithelial cells in the scRNA-Seq dataset (**Figure 2B**). Integration scores, a reflection of confidence in the assignment of cell identity, were highest among terminally differentiated cell types (e.g., exocrine and hormone-expressing endocrine cells) (**Figure S2A**), indicating less ambiguity in chromatin accessibility once cell fate is determined. Even in the absence of integration with scRNA-Seq data, all cell types were identified when clustering on chromatin accessibility alone (**Figure S2B**).

**Figure 2:**
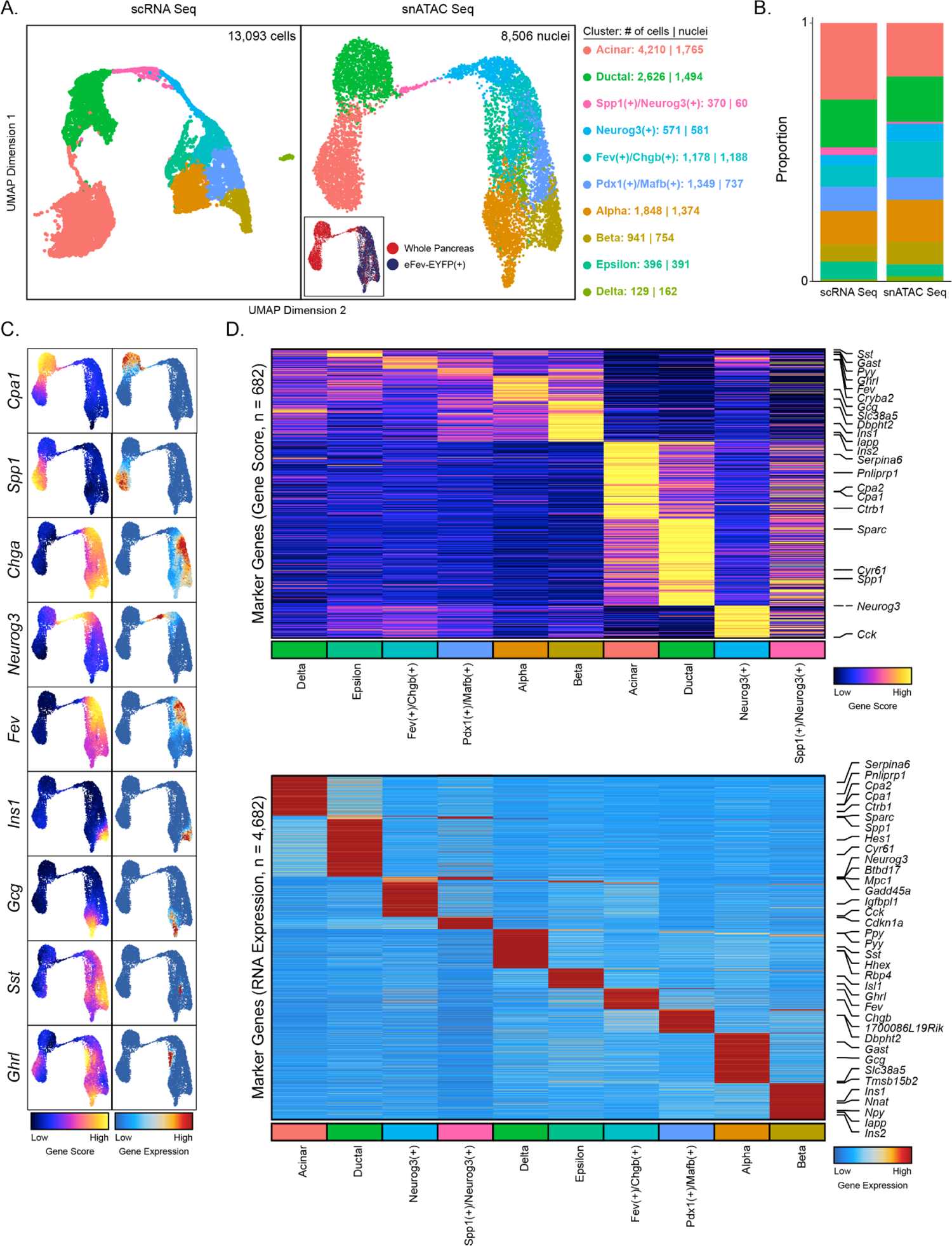
Integration of single-cell transcriptional and chromatin accessibility data identifies epithelial heterogeneity in the developing murine pancreas. (**A**) UMAP plots enabling visualization of scRNA-Seq (left) and snATAC-Seq (right) data for all epithelial cells in the E14.5 pancreas. Numbers of cells/nuclei are depicted on the right, along with cell type annotations. The scRNA-Seq dataset was previously published by our group [19]. (**B**) Bar graph depicting the proportion of all each cell type as a fraction of all epithelial cells in the scRNA-Seq and snATAC-Seq datasets. Colors match the cell types in (**A**). (**C**) Feature plots showing chromatin accessibility (Gene Score; left) or Gene Expression (right) of genes that mark each epithelial cell type. *Cpa1, Spp1, Chga*, *Neurog3*, *Fev*, *Ins1*, *Gcg*, *Sst*, and *Ghrl* mark acinar, ductal, pan-differentiated endocrine, early endocrine progenitor (EP), intermediate EP, beta, alpha, delta, and epsilon cells, respectively. (**D**) Heatmaps depicting genes that are differentially accessible (gene score; top heatmap) or differentially expressed (gene integration matrix; bottom heatmap) across the epithelial clusters. Genes listed were selected from the set of genes determined to be differentially expressed among scRNA-Seq clusters in panel (**A**).

Next, we confirmed the cell type annotations by assessing chromatin accessibility (gene score), as well as the transferred RNA expression from the integration (gene expression). We observed high concordance between chromatin accessibility and RNA expression of the marker genes defining our cell types (**Figure 2C**). Additionally, we observed cell-type specific chromatin accessibility of each marker gene locus (**Figure S2C**). When assaying differentially-accessible or -expressed genes, we observed far fewer significantly differentially accessible genes (n = 682) compared to differentially expressed (n = 4,682) (**Figure 2D, Supplemental table 1**). Among these differentially-accessible genes were top markers of each cluster identified by differential gene expression analysis of our scRNA-Seq dataset. Taken together, these data confirm the existence of heterogeneous epithelial populations initially identified by scRNA-Seq, here by an orthogonal method.

### 3.3. Identification of candidate regulators of epithelial cell fate

To identify regulators of cell fate decisions in the developing pancreatic epithelium, we applied the peak calling algorithm MACS2 [46] to our dataset. We identified 169,197 peaks across all epithelial clusters, with 32,710 peaks exhibiting differential accessibility across cell types (**Figure 3A, Supplemental table 2**). Next, we assayed for TF motif enrichment in these differential peaks, identifying 348 enriched motifs. A number of TFs in the same family were deemed enriched due to the similarities in DNA binding motifs. For instance, TFs with enriched motifs included known regulators of pancreatic epithelial development, such as Sox family members (*Sox2, Sox4, Sox9*; Ductal), Hox family members (*Hoxb4, Hoxc4, Hoxa4*; Beta) and members of the Rfx family (*Rxf3* through *Rfx7*; Fev(+)/Chgb(+) and Pdx(+)/Mafb(+)) (**Supplemental table 2**). To distinguish among TFs with similar DNA motifs identified in a given cell type, we next identified significant TF motif deviations (calculated as deviation of motif enrichment in accessible peaks from the expected distribution based on the average across all cells) of each cell type using ChromVAR [40]. The TFs from ChromVAR were then correlated with their gene expression profiles from the integrated RNA expression matrix, thereby identifying so-called “correlated TFs” that are both expressed and have significant motif deviation (**Figure 3B, Supplemental table 3**). By breaking this down further on a per-cluster basis, we then were able to observe the cell type-specific motif deviations and gene expression of the correlated TFs, narrowing the number of TFs with enriched motifs from 348 (**Figure 3A**) to 48 correlated TFs (**Figure 3B, C**). Correlated TFs included multiple members of the Fox family (*Foxo1*, Delta cells; *Foxj2* and *Foxc1*, Epsilon cells; *Foxp2* and *Foxp3*, Alpha cells; *Foxp1*, Fev(+)/Chgb(+) cells), as well as the Sox family (**Figure 3C**). Interestingly, by observing not only motif deviation but also gene expression, we were able to determine that although both Ductal cells and Spp1(+)/Neurog3(+) EPs showed high motif deviation of Sox4, expression was significantly higher in the latter population (**Figure 3D**). This is in line with previously published work that shows that *Sox4* works with *Neurog3* to induce endocrine differentiation in the developing murine pancreas [47].

**Figure 3:**
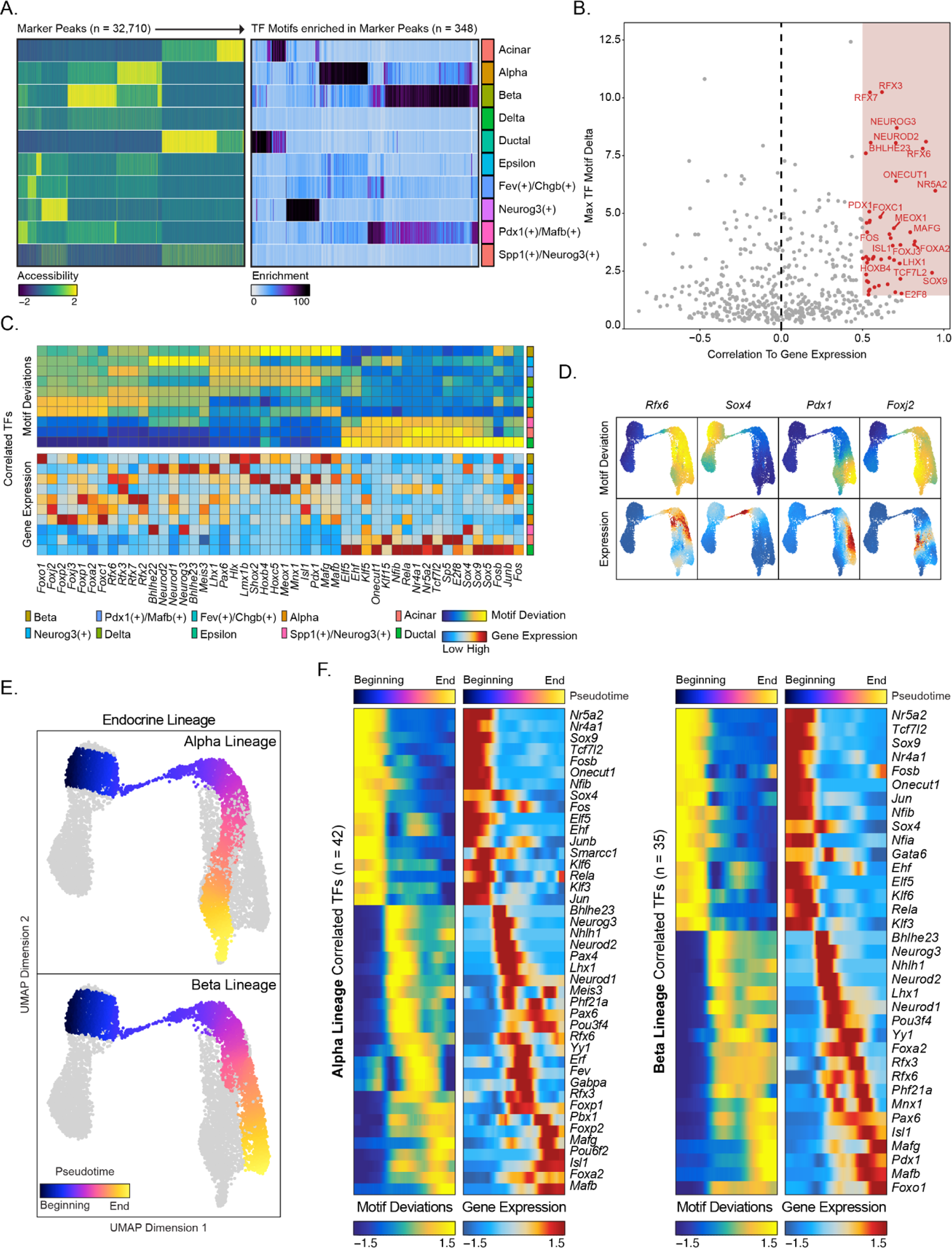
Identification of candidate correlated transcription factors governing pancreatic epithelial cell fate. **(A)** Heatmaps depicting enriched marker peaks (left) and transcription factor (TF) motifs enriched in marker peaks (right) for each epithelial cell type. (**B**) Dot plot shows so-called “correlated” TFs (those with high correlation between motif deviation score and gene expression) in all epithelial cells. (**C**) Heatmaps revealing cell type-specific motif deviation scores (top) and gene expression values (bottom) of positive TFs identified in (**B**). (**D**) Feature plots displaying motif deviation (top) and gene expression (bottom) of selected positively correlated TFs at single-cell resolution. (**E**) Pseudotemporal ordering of epithelial cells along the Alpha and Beta lineages based on chromatin accessibility. (**F**) Heatmaps depicting positively correlated TFs across pseudotime (from left to right) for Alpha (left heatmaps) and Beta (right heatmaps) lineages.

We next sought to understand the correlated TFs across the Alpha and Beta cell lineages. We first calculated the pseudotime values of cells along both trajectories (**Figure 3E**) and then applied the same motif deviation and gene expression correlation analysis for the genes and enriched motifs along these lineages (**Figure 3F, Supplemental table 3**). Across the Alpha lineage (including Ductal, Spp1(+)/Neurog3(+), Neurog3(+), Fev(+)/Chgb(+), and Alpha cells) we identified 42 correlated TFs. This included TFs in Ductal cells (*Nr5ac, Nr4a1, Sox9*), progenitor cells (*Sox4, Neurog3, Neurod2, Pax4*), and Alpha cells (*Foxp2, Isl1, Mafb*). For the Beta lineage, we identified 35 correlated TFs, including *Mnx1, Mafg, Pdx1*, and *Foxo1*. In summary, the multi-layered approach taken here has further distilled the subset of TFs that likely play an important role in governing fate selection during endocrinogenesis.

### 3.4. Gene regulatory networks controlling epithelial cell fate

Our analyses thus far have identified accessible chromatin and correlated TFs within the epithelial compartment of the developing endocrine pancreas. How and where these TFs bind and affect downstream target genes to govern cell fate decisions is not as well understood, however. To address this gap in knowledge, we next sought to construct a gene regulatory network (GRN) for Acinar, Ductal, and endocrine cells of the Alpha and Beta lineages (**Figure 4A**). We utilized the computational pipeline Integrated Regulatory Network Analysis (IReNA) v2 [48] (**Figure 4B, Figure S4A**), which combines both scRNA-Seq and snATAC-Seq data to predict TF binding of downstream target genes in a cell type-specific manner. First, we performed differential gene expression analysis on our scRNA-Seq dataset to identify genes enriched in each cell type (**Figure S4B, Supplemental table 4**). We then performed peak-to-gene linkage analysis in our integrated scRNA- and snATAC-Seq datasets, identifying accessible regions of chromatin (peaks) that are either positively or negatively significantly correlated with gene expression (genes) (**Figure S4C**). These peak-to-gene peaks were then further filtered and annotated as correlated accessible regions (CARs) belonging to one of three categories: TSS (when the peak lies in the transcription start site (TSS) for the gene), gene body (when the distance between peak and TSS is less than 100 kb, and the peak-to-gene score calculated above is significant), or distal (when the peak is 100 kb upstream or downstream of the TSS of a correlated gene, and the peak-to-gene score is significant). We observed varying proportions of CAR categories among the cell types, with TSS representing the highest proportion, followed by positive and then negative in the Ductal, Acinar, and early EP populations (**Figure S4D**). As endocrine differentiation progresses, the majority of CARs are positively correlated with expressed genes. We next predicted the cell-type specific TF binding in these CARs by searching for TF DNA binding motifs in the CARs. Once identified, we then filtered the TFs by calculating their TF footprint score, retaining TFs with a score deemed significant by IReNA.

**Figure 4:**
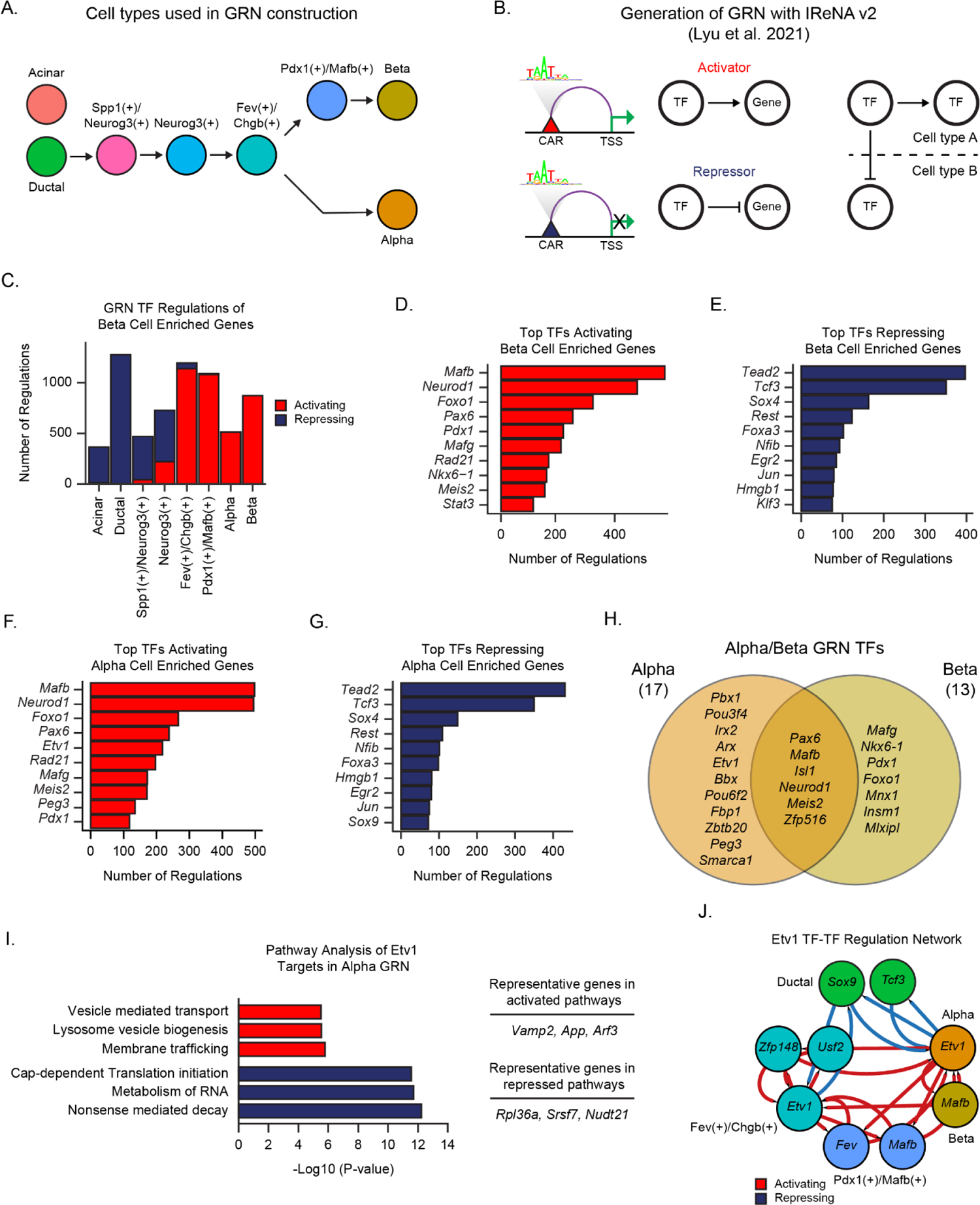
Mapping the gene regulatory networks active in the Alpha and Beta cell populations. (**A**) Schematic depicting application of the Integrated Regulatory Network Analysis (IReNA) v2 pipeline to identify gene regulatory networks (GRNs) within specific cell types. The GRN refers to active, cell type-specific transcription factors (TFs) and their target genes. (**B**) Cell populations used as input for IReNA. Arrows denote known lineage relationships. (**C**) Bar graph indicating the number of downstream target genes for TFs enriched in the Beta cell type. Activating (red bar) and repressing (blue bar) refers to positive or negative correlation between gene expression levels of the TF and the target gene. (**D-E**) Bar graphs showing the top 10 TFs with the highest number of activating (**D**) or repressing (**E**) regulations of target genes enriched in the Beta cell type. (**F-G**) Bar graphs showing the top 10 TFs with the highest number of activating (**F**) or repressing (**G**) regulations of target genes enriched in the Alpha cell type. (**H**) Venn diagram depicting the overlap between Alpha and Beta GRN TFs. (**I**) Bar graph depicting top most significantly (p-value < 0.01) enriched pathways of genes activated (red bars) or repressed (blue bars) by the TF *Etv1* in the Alpha cell GRN. (**J**) Network diagram representing regulations between *Etv1* and interacting TFs. Each TF is represented by a circle (node) that is colored by the cell type in which that TF is active in the GRN. Activating regulations are depicted by red lines, while repressing are depicted by blue lines.

We observed the highest number of GRN TFs in the Fev(+)/Chgb(+) population (39 TFs), followed by Neurog3(+) (33 TFs), and Ductal (29 TFs) cells (**Figure S4E, Supplemental Table 5**). Among our network of TFs and target gene-associated CARs, we binned these interactions as either activating or repressing by correlating the expression of each TF-target gene pair. Genes with a positive TF-target gene correlation were annotated as being activated by their given TF, while those with a negative TF-target gene correlation were annotated as being repressed. The Ductal and the Fev(+)/Chgb(+) populations had the highest number of regulations, followed by Pdx1(+)/Mafb(+) and Neurog3(+) (**Figure S4F**). The regulations among all the populations examined were relatively evenly split between activating and repressing. Lastly, from the GRN constructed above, we identified pairs of TFs that regulated one another; for each TF, we identified target genes that are also TFs and mapped these pairs as either activating or repressing depending on the correlation of gene expression of the target TF in the given cell type. This analysis permitted us to identify TFs that regulate the expression of other TFs in given cell types (**Figure S4G, Supplemental table 6**).

To examine the TFs comprising this epithelial GRN in more depth, we first focused on the hormone(+) populations within our dataset. We found that genes enriched in the Beta cell population are largely repressed in the Acinar and Ductal GRNs, then gradually become activated as endocrine differentiation proceeds (**Figure 4C**). Within the Alpha cell population, Beta cell enriched genes are almost exclusively activated, consistent with previous studies investigating gene expression of individual TFs revealing that beta and alpha cells share common expression of genes needed for proper development and function [7]. We then inquired within all of the GRNs defined for epithelial cell types, which TFs either activate or repress genes enriched in the Beta cell population. Among the top activating TFs were known regulators of Beta cell development, such as *Mafb, Neurod1, Pdx1*, and *Nkx6-1* (**Figure 4D**). Targets of these activating TFs identified by our GRN analysis included numerous genes, both known and novel (**Supplemental table 5**). Repressors of Beta cell enriched genes are largely contained within the Ductal GRN and include TFs such as *Tead2, Sox4,* and *Rest* (**Figure 4E**). Conversely, TFs activating Alpha cell enriched genes include Beta cell activating TFs such as *Mafb* and *Neurod1* (**Figure 4F**). TFs repressing Alpha cell enriched genes also included *Tead2, Sox4,* and *Rest* (**Figure 4G**), suggesting that these TFs repress global hormone(+) cell gene signatures. TFs that overlapped between the Alpha (17 TFs) and Beta (13 TFs) GRNs comprised known endocrine regulators, such as *Pax6, Mafb, Neurod1,* and *Isl1*, as well as TFs less well studied in pancreas, such as *Zfp516* and *Meis2* (**Figure 4H**). TFs unique to the Beta GRN included known regulators of beta cell fate, such as *Nkx6-1*, *Pdx1,* and *Foxo1*, while those less well characterized included *Mlxipl*. Examples of TFs unique to the Alpha GRN were *Irx2* and *Arx*, known regulators of Alpha cell fate. Less well characterized TFs included *Pbx1*, which is required for proper pancreas development [49], *Bbx, Peg3,* and *Etv1*.

Among the top activating TFs of Alpha cell enriched genes was *Etv1* (**Figure 4F**), reported to be a direct or indirect target of *Nkx2-2* [50]. Within beta cells, failure to properly degrade *Etv1*, *Etv4*, and *Etv5* results in impaired insulin secretion [51]. We took a closer look at the downstream targets of *Etv1* in the Alpha cell GRN and found that *Etv1* was predicted to activate 140 genes, and repress 133 genes, in the Alpha GRN (**Supplemental table 5**). When performing pathway analysis on these genes, we observed that *Etv1* activated pathways related to vesicle mediated transport, lysosome vesicle biogenesis, and membrane trafficking (**Figure 4I**). Pathways repressed by *Etv1* included translation initiation, metabolism of RNA, and nonsense-mediated decay. When examining TF-TF interactions, we found that *Etv1* is repressed by *Sox9* and *Tcf3* in the Ductal population, and activated by *Mafb* and *Fev* in the Pdx1(+)/Mafb(+) population and by *Usf2* and *Zfp148* in the Fev(+)/Chgb(+) population (**Figure 4J**).

In summary, the computational analyses described in this section have permitted the construction of a gene regulatory network of the acinar, ductal, and major endocrine lineages in the developing mouse pancreas. This workflow, which is dependent on integration of both chromatin accessibility and transcriptional data, has identified regulators of alpha and beta cell fate that can serve as the subjects of further experimental study.

### 3.5. Gene regulatory networks governing the initiation of endocrine differentiation

During mammalian pancreatic development, a subset of cells within the branching ductal epithelium activate the expression of the master regulator of endocrine differentiation, *Neurog3*. These rare Neurog3(+) cells represent the earliest known EP population, and considerable attention has been devoted to understanding the ductal to EP transition. In addition to *Neurog3*, which is required for mouse endocrine differentiation [6], numerous other TFs have been identified that are also important for endocrinogenesis. Investigation of *NEUROG3* binding across the genome in human pluripotent stem cell (hPSC)-derived EP cells revealed widespread regulation of 138 TFs, some with known roles in endocrine development and others with unknown function [52]. Further studies in human cells used inducible and knockout models in hPSC-derived endocrine cells to identify predicted targets of multiple endocrine TFs, including *NEUROG3*, *PDX1,* and *RFX6* [53]. Generation of an Ngn3-timer fluorescent reporter mouse line that permitted the specific isolation of early Ngn3-expressing cells identified numerous putative direct targets of *Neurog3* in mouse EPs [54]. These studies have highlighted the need for a broad, integrated analysis of all TFs and downstream targets that control the initiation of endocrine differentiation.

Towards this end, we began by investigating the GRN regulating the transition from a ductal to EP cell state. TFs in the Ductal GRN promoting the expression of Spp1(+)/Neurog3(+) EP genes include known regulators of endocrine cell fate, such as *Sox4* and *Sox9* [47, 55] (**Figure 5A**). Interestingly, *Tead2* and *Tcf3* activated the most genes enriched in the Spp1(+)/Neurog3(+) and Neurog3(+) EP populations (**Figure 5B**), indicating that these TFs are important initiators of an endocrine cell fate. The Yap/Tead signaling complex has previously been shown to activate multipotent progenitor cell enhancers and regulate epithelial outgrowth during human pancreatic development [56]. *Tcf3*, also known as *E47*, has been shown in a human cell line to dimerize with *NEUROG3* to bind to the promoter region of the *INSM1* gene [57], which is required in mice to maintain mature beta cell function [58]. TFs involved in the transition from a Spp1(+)/Neurog3(+) to a Neurog3(+) EP cell state include well known regulators of endocrine differentiation, such as *Nkx2-2*, *Pax4*, *Neurod2*, and *Neurog3* (**Figure 5A**). Major repressors of EP enriched genes in the Ductal GRN include TFs such as *Rest* and *Nfib* (**Figure 5C**). *Rest* is a master regulator of neurogenesis and has been previously described to inhibit direct reprogramming of pancreatic exocrine to endocrine cells by inhibiting the binding of *Pdx1* to key endocrine differentiation-related genes [59]. In addition, loss of *Rest* results in increased generation of pancreatic endocrine cells during development [60, 61]. *Nfib* belongs to the Nuclear Factor I protein family, of which another member, *Nfia*, plays a role in the induction of EP cell fate [62]. Among TF-TF pairs identified in Ductal, Spp1(+)/Neurog3(+), and Neurog3(+) cells, most were classified as activating, with the exception of *Hes1, Fos* (Ductal), and *Tcf3* (Spp1(+)/Neurog3(+)) (**Figure S5A**). The classification of *Hes1* as repressing is consistent with what is known about the role of Notch signaling in the initiation of EP cell fate [5, 63]. Taken together, our GRN analysis has identified novel candidate regulators, such as *Tcf3* and *Tead2*, of the ductal to EP cell state transition. These results expand upon our knowledge of this key developmental transition and serve as a resource for future studies.

**Figure 5:**
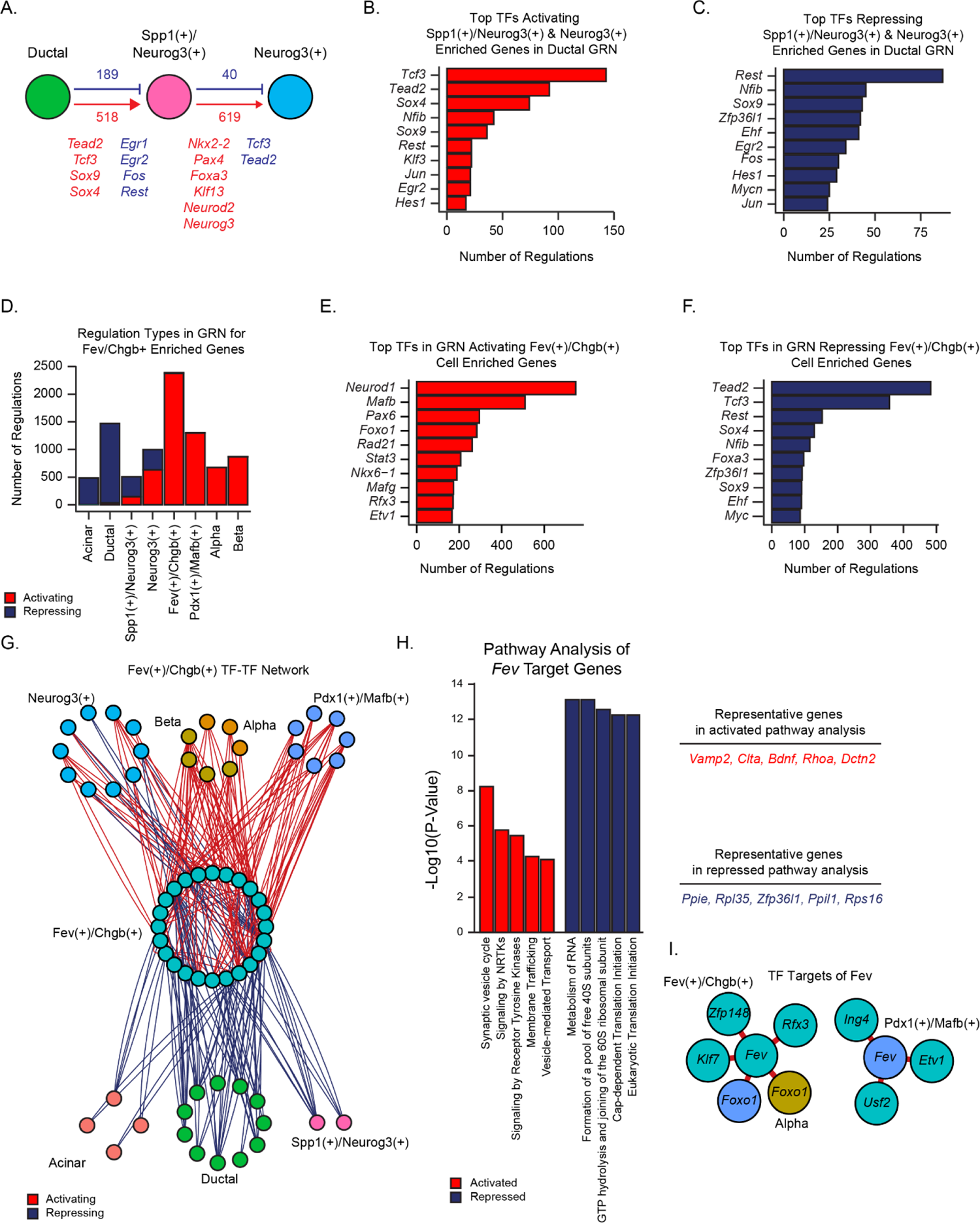
Mapping the gene regulatory networks active in the Fev-expressing pancreatic endocrine population. (**A**) Cartoon depicting the number of activating (red) and repressing (blue) regulations between transcription factors (TFs) in Ductal, Spp1(+)/Neurog3(+) and Neurog3(+) populations, with representative TFs indicated. (**B-C**) Bar graphs depicting the top 10 TFs in the Ductal GRN with the highest number of activating (**B**) or repressing (**C**) regulations of target genes that are enriched in the Spp1(+)/Neurog3(+) and Neurog3(+) cell types. (**D**) Bar graph depicting the number of TF regulations of target genes enriched in the Fev(+)/Chgb(+) cell type. Activating (red bars) and repressing (blue bars) refers to positive or negative correlation, respectively, of gene expression between the TF and target gene. (**E-F**) Bar graphs depicting the top 10 transcription factors with the highest number of activating (**E**) or repressing (**F**) interactions of target genes enriched in the Fev(+)/Chgb(+) cell type. (**G**) Network diagram depicting all TF-TF regulations between TFs enriched in the Fev(+)/Chgb(+) GRN and all other GRNs. TFs are denoted by each node, which is colored by the cell type in which the TF is found. Each activating regulation is depicted by a red line, while each repressing regulation is depicted by a blue line. (**H**) Bar graph depicting top significant (p-value < 0.01) pathways of genes activated (red bars) or repressed (blue bars) by *Fev* in the GRN analysis. (**I**) Network diagram depicting the TFs activated (red lines) by *Fev* in the GRN analysis. Each node (TF) is colored according to the cell type in which it is expressed.

We next focused on Fev(+)/Chgb(+) cells, as our previous work indicated that this cell state represents the bifurcation point at which the Alpha or Beta lineage is established (**Figure 4A)** [19]. As expected, we observed that the Acinar and Ductal cell types largely repress genes that are enriched in the Fev(+)/Chgb(+) population (**Figure 5D**). These genes begin to be activated as an endocrine cell fate is established (Spp1(+)/Neurog3(+) and Neurog3(+) cell types) and are fully activated by the Fev(+)/Chgb(+) cell stage. Curiously, the Fev(+)/Chgb(+) enriched genes are not repressed in the Alpha and Beta cell types, suggesting that Alpha/Beta cell fate is due more to activation of key Alpha/Beta genes as opposed to the repression of progenitor-associated genes. Among the top TF activators of Fev(+)/Chgb(+) enriched genes (**Supplemental table 5**), we observed that known regulators of endocrine cell fate such as *Neurod1, Mafb*, and *Pax6* activated the most genes (**Figure 5E**). Activators also included less well described TFs, such as members of the Rfx family (*Rfx3, Rfx6*), as well as *Foxo1* and *Etv1* (**Figure 5E**; **Supplemental table 5**). Conversely, TF repressors of Fev(+)/Chgb(+) enriched genes included the TFs *Tead2*, *Tcf3*, *Rest, Sox4*, *Sox9*, and *Nfib,* among others (**Figure 5F**). Next, we constructed a network diagram of TF pairs that either activate or repress TFs enriched in the Fev(+)/Chgb(+) population (**Figure 5G**). Consistent with our observations in **Figure 5D**, TF-TF regulations in the Acinar, Ductal, and Spp1(+)/Neurog3(+) cell states were entirely repressive, and regulations in the Neurog3(+) cell state were a mix of activating and repressing (**Figure 5G; Supplemental table 6**). In contrast, TF-TF regulations in the Alpha, Beta, and Pdx1(+)/Mafb(+) states were exclusively activating (**Figure 5G; Supplemental table 6**).

When comparing the GRNs among all progenitors and precursors, we identified 18 TFs unique to the Fev(+)/Chgb(+) population (**Figure S5B, Supplemental table 6**). Among these TFs identified within the Fev(+)/Chgb(+) GRN, *Mafb* was the top activator of the transition from Fev(+)/Chgb(+) to either Pdx1(+)/Mafb(+), Alpha, or Beta cell states (**Figure S5C-E**). Among the top 10 TFs with the highest number of activating regulations across the transition from a Fev(+)/Chgb(+) to Pdx1(+)/Mafb(+) cell state was Vitamin D receptor (*Vdr*), whose expression has been linked to beta cell function and diabetes (**Figure S5C**) [64]. As expected from **Figure 4F**, *Etv1* had a higher number of activating regulations for Alpha cell enriched genes compared to Pdx1(+)/Mafb(+) or Beta populations (**Figure S5C-E**), and *Pax6* was identified as one of the top TFs activating beta cell enriched genes (**Figure S5E**). Top 10 TFs activating Alpha cell enriched genes included *Foxp1*, which has been implicated in postnatal alpha cell expansion and function (**Figure S5D**) [65].

The gene *Fev* was initially described as a prototypical serotonergic transcription factor in the brain [66], then as a gene expressed in developing and adult mouse pancreatic islets [67]. More recently, we found that *Fev* in the pancreas marks an intermediate progenitor of the mouse endocrine lineage [19]. In a beta cell line, *Fev* has been demonstrated to bind not only to serotonergic genes, reflective of common transcriptional cascades that drive the differentiation of both serotonergic neurons and of beta cells [68], but also to a conserved insulin gene regulatory element [67]. While levels of glucagon, somatostatin, pancreatic polypeptide, or ghrelin were unchanged in Fev whole body knockout (Fev-/-) embryos, levels of *Ins1*, *Ins2*, and islet amyloid polypeptide (*Iapp*) were reduced, as were the levels of the gene *Slc2a2* (which encodes Glut2). Ohta et al. also measured the expression of multiple genes encoding islet TFs in Fev-/- embryos, and found that loss of Fev did not alter the expression of *Isl1*, *Mafa*, *Mafb*, *Mnx1*, *Neurod1*, *Pax6*, *Pdx1*, *Rfx6*, *Insm1*, *Nkx2-2,* or *Nkx6-1* at E18.5. Knockout embryos did display slightly lower levels of *Lmx1b*, which also plays a role in serotonergic neuron development downstream of *Nkx2-2*.

Our GRN analysis computed 110 genes activated and 69 genes repressed by *Fev* (**Supplemental table 5**). Pathway analysis of activated genes included pathways such as synaptic vesicle cycle, signaling by NRTKs, and vesicle mediated transport (**Figure 5H**). These data corroborate previous findings in which full body knockout of *Fev* resulted in decreased pancreatic insulin content, an impairment of insulin secretion, and consequently defects in glucose clearance [67]. Pathway analysis of repressed genes included many translation-associated pathways, such as metabolism of RNA, cap-dependent translation initiation, and formation of a pool of free 40s subunits (**Figure 5H**). Downstream TF interactions of *Fev* were all activating and included the TFs *Rfx3*, *Klf7*, and *Foxo1* (**Figure 5I**).

Taken together, our data identify both known and novel regulators of pro-Alpha and pro-Beta cell fates that are active in the Fev(+)/Chgb(+) stage, the cell state that represents the bifurcation point in the endocrine differentiation trajectory. Our analysis also yields a comprehensive view of potential targets of *Fev,* as well as insights regarding its function in regulating the machinery required for the production of endocrine hormone-containing vesicles.

### 3.6. Characterization of chromatin accessibility and identification of GRNs within pancreatic mesenchymal cell types

Although proper development of the pancreatic epithelium depends on signals from the surrounding mesenchyme, the lineage and function of pancreatic mesenchymal cells remains vastly understudied. In previous work, we and others have used scRNA-seq to identify multiple transcriptionally distinct mesenchymal populations, including mesothelium, within the developing murine pancreas [19, 20]. Still, the upstream genetic regulators that maintain these distinct cell states are not defined. Data from snATAC-Seq of pancreatic mesenchymal cells would shed light on whether distinct states of chromatin accessibility correspond to transcriptionally distinct cell subpopulations and would reveal which TFs and binding sites are actively involved in controlling mesenchymal cell state.

We integrated the snATAC-Seq data from the mesenchymal populations within our dataset with the age-matched (E14.5) scRNA-Seq data, using methods as described above. Clustering of the scRNA-Seq dataset identified six populations of mesenchymal cells, including one cluster enriched in the expression of *Gap43* (Gap43(+)), another cluster enriched in expression of *Sfrp2* (Sfrp2(+)), two clusters expressing chemokines (Cxcl12(+) and Cxcl13(+)), Vascular Smooth Muscle cells (VSM; Acta2(+)) and finally Mesothelium (Wt1+) (**Figure 6A**). Integration and cell label transfer classified all populations in our snATAC-Seq dataset, with the exception of the small Cxcl13(+) population (**Figure 6A,B**). Clustering on chromatin accessibility alone, without integration with scRNA-Seq data, still resulted in the identification of similar clusters as in the integrated dataset (**Figure S6A**).

**Figure 6:**
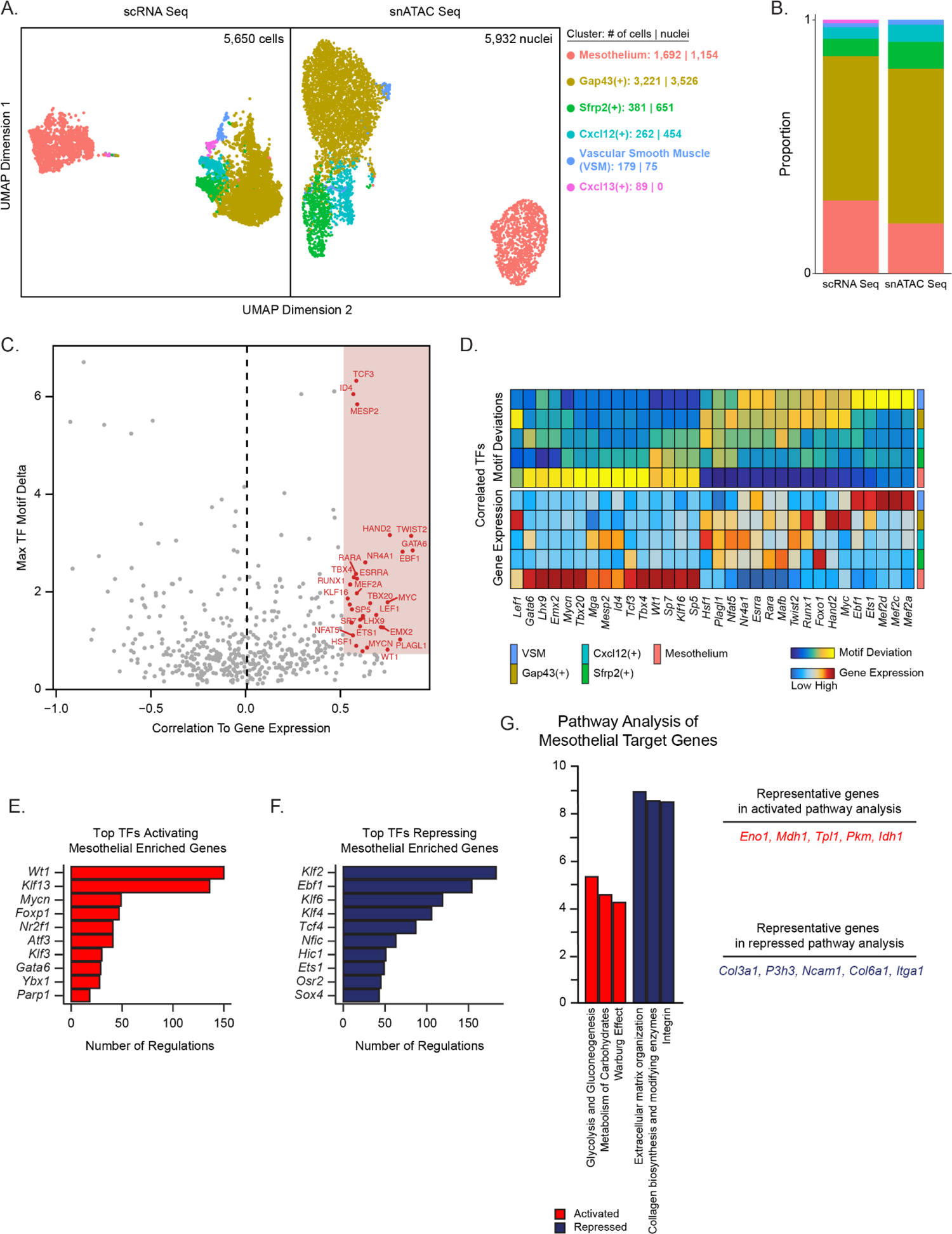
Chromatin accessibility and gene regulatory network in the developing pancreatic mesenchyme. (**A**) UMAP plots enable visualization of scRNA-Seq (left) and snATAC-Seq (right) data for all mesenchymal cells in the E14.5 pancreas. Numbers of cells/nuclei are depicted on the right, along with cell type annotations. The scRNA-Seq dataset is from our previously published work [19]. (**B**) Bar graph depicts the proportion of each cell type in the scRNA-Seq and snATAC-Seq datasets. Colors match the cell types in (**A**). (**C**) Dot plot showing correlated transcription factors (as determined by correlation between motif deviation score and gene expression) in mesenchymal and mesothelial cells. (**D**) Heatmaps reveal cell type-specific motif deviation scores (top) and gene expression values (bottom) of positive transcription factors identified in (**C**). (**E, F**) Bar graph depicting the top TFs activating (**E**) and repressing (**F**) genes enriched in the mesothelial population. (**G**) Bar graph depicting top significant (p-value < 0.01) pathways of genes activated (red bars) or repressed (blue bars) by mesothelial cells in the GRN analysis. VSM, vascular smooth muscle.

Next, we identified differentially accessible peaks across all of the mesenchymal populations and found a total of 109,660 peaks, with 30,737 peaks displaying differential accessibility (**Figure S6B, Supplemental table 2**). The majority of these differentially accessible peaks were enriched in the Gap43(+) and Mesothelial populations, with the Cxcl12(+) and Sfrp2(+) populations showing a more modest number. Motif enrichment in these differentially accessible peaks identified 123 enriched motifs (**Figure S6B**). Correlation of gene expression and motif deviation scores identified 32 correlated TFs (**Figure 6C,D**), including known regulators of mesenchymal cell fate, such as *Wt1*, *Twist2*, and *Hand2*.

We next assembled the mesenchymal GRN with IReNA v2 as described above. Among the mesenchymal cell types, the Gap43(+) and Mesothelial populations had the highest number of total regulations (**Figure S6C**) as well as the highest numbers of TFs in each of the GRNs (**Figure S6D**). In the TF-TF network, TFs active in the GRN of Mesothelium largely repressed the expression of Gap43(+)-associated TFs, whereas the Gap43(+) population activated Cxcl12(+)-associated TFs (**Figure S6E**). We next assessed which GRN TFs were either exclusive or shared among the mesenchymal populations (**Figure S6F**). The GRN in the Gap43(+) population contained 18 specific TFs, including genes involved in epithelial-mesenchymal transition (EMT) such as *Snai2, Twist2,* and *Zeb1*, suggesting that cells within the Gap43(+) population are actively undergoing EMT processes during pancreatic development. Other cell type-specific TFs included *Nkx2-3*, *Hoxb4,* and *Prxx1*. TFs exclusive to the Cxcl12(+) population included members of the nuclear factor 1 (*Nfia* and *Nfic*), Ap-1 (*Junb* and *Jund*), and Klf families (*Klf3*, *Klf6; Klf2* shared with the Gap43+ population). TFs specific to the Sfrp2(+) GRN included *Zfhx3, Hoxb5,* and *Meis2,* while *Pbx1, Tcf21,* and *Bcl11a* were shared with the Cxcl12(+) population, and *Barx1* was shared with the Mesothelial population.

We next focused on mesothelial cells, which consist of a monolayer of specialized cells that line the pleura and internal organs of adult tissues and serve numerous functions in the adult, including lubrication of tissue and immune surveillance [69, 70]. In the developing lung, lineage tracing studies have demonstrated that mesothelium also acts as a progenitor for certain specialized mesenchymal cell subtypes [71]. Furthermore, previous work in our lab predicted the downstream lineages of mesothelial cells in the developing pancreas based on pseudotemporal ordering of scRNA-Seq data [19]. Despite this, relatively little is known about how mesothelial cells are formed and maintained during pancreatic development. Within the snATAC-Seq dataset, we identified 23 TFs uniquely active within the GRN of the Mesothelial population (**Figure S6F**). Top TFs activating Mesothelial-enriched genes included *Wt1,* which has previously been shown to be a master regulator of mesothelial formation [72], along with *Klf13,* whose role in mesothelial cell development and homeostasis is not well understood and thus warrants further study (**Figure 6E**). Top repressing TFs of Mesothelial-enriched genes included *Klf2* (active in Gap43(+) and Cxcl12(+) cells, **Figure S6F**), *Ebf1* (active in Gap43(+), VSM, and Cxcl12(+) cells) and *Klf6* (active in Cxcl12(+) cells) (**Figure 6F**). Pathway analysis of activated and repressed genes in the Mesothelium GRN identified pathways related to metabolism as activated, and ECM formation and deposition as repressed (**Figure 6G**).

Taken together, profiling of the chromatin accessibility within the cells of the developing mouse pancreatic mesenchyme has determined differentially accessible peaks among these populations, and identified TFs potentially important in mesenchymal development through the use of motif enrichment and gene expression correlation analyses. Lastly, we have constructed a set of mesenchymal GRNs, identifying active TFs and their downstream target genes. These data will provide a resource for future work geared towards studying mesenchymal biology and understanding how this important but understudied non-epithelial population is maintained.

## 4. Discussion

Numerous studies have used scRNA-Seq to characterize developing mouse pancreas tissue, providing important insights into cellular heterogeneity and key transcriptional programs expressed in developing cell types [19–23]. Still, these datasets lack information about which of the expressed TFs are active and binding, and about how the TFs are organized into regulatory networks. Profiling of the chromatin accessibility landscape at single-cell resolution has emerged as a powerful approach for generating new insights about regulatory programs governing development and cell fate decisions across multiple tissue types [31,32,73,74], and we have now extended this approach to developing mouse pancreas tissue. Given that we were particularly interested in interrogating mechanisms underlying endocrine lineage allocation, we utilized a genetic tool to significantly enrich for EP cells. Previous work from our laboratory had identified the transcription factor *Fev* as a marker of a novel endocrine progenitor state, and lineage reconstruction analysis indicated that it is at this *Fev*(+) state that lineage allocation is executed [19]. Here, we validated the use of an eFev-EYFP transgenic mouse line for enriching for *Fev*-expressing endocrine cells in the developing pancreas.

We have generated a comprehensive atlas of chromatin accessibility in the developing E14.5 murine pancreas, including enriched endocrine populations as well as non-endocrine cell types. Although previous studies have investigated chromatin accessibility in the developing pancreatic epithelium through bulk ATAC-Seq of sorted populations [20, 26], to our knowledge our study represents the first to examine chromatin accessibility at true single-cell resolution. By integrating both scRNA-Seq and snATAC-Seq data, we successfully generated a refined list of correlated TFs that are not only expressed, but also likely binding to open regions of chromatin to control cell fate decisions. Furthermore, we constructed cell-type specific GRNs describing active TFs and their putative target genes through the binding of *cis*-regulatory regions. Our analysis identified a number of known regulators of endocrine cell fate, such as TFs *Pdx1* and *Nkx6-1* in beta cells and *Arx* in alpha cells, as well as identified novel candidate TFs, such as *Mlxipl* in beta cells and *Pbx1* and *Peg3* in alpha cells. Although here our focus within the epithelial compartment was on the endocrine lineages, our dataset also provides a rich resource for future interrogation of gene regulatory networks controlling acinar and ductal cell fates. Identification of these networks will inform efforts underway at generating stem cell-derived exocrine cells *in vitro* for studies aimed at understanding exocrine cell physiology and modeling of diseases such as cystic fibrosis, pancreatitis, and pancreatic cancer [75, 76].

Traditional single-gene studies, along with genomic studies, have led to the identification of numerous TFs that play a functional role in regulating pancreatic endocrine differentiation. Although some individual TF-TF interaction pairs have been identified through these methods, the field still lacks an understanding of how these TFs are broadly arranged in regulatory networks across cell types and developmental stages. Our analysis permitted the creation of a TF-TF regulatory network, identifying TFs that control cell fate decisions through the binding and regulation of other important TFs. The assembly of these networks has identified both known and novel TF-TF interacting pairs whose associations can be experimentally validated in future studies using tools such as Chromatin Immunoprecipitation Sequencing (ChIP-Seq) for confirmation of binding to specific DNA regions. Furthermore, CRISPR-mediated gene editing can be used to assess the downstream consequences of loss of individual candidate TFs on cell fate outcomes.

In contrast to the pancreatic epithelium, the cellular composition and transcriptional features of the pancreatic mesenchyme have been less well described. We and others have applied scRNA-Seq to mesenchymal tissue to identify transcriptionally distinct sub-populations [19, 20] and infer lineage relationships among some of these cell subtypes [19]. In addition, functional heterogeneity among pancreatic mesenchymal cells has begun to be explored. For instance, one study reported that expression of *Pbx1* in a subset of Nkx2-5+ mesenchymal cells defines an anatomically specialized, pro-endocrine niche [24]. Which genes, including TFs, govern the acquisition of mesenchymal subpopulation identity, however, is poorly understood. Our work begins to investigate novel TFs regulating mesenchymal cell fate and will serve as an important resource for understanding mesenchymal development and function. Our dataset identifies which gene networks should be activated in order to generate not only organ-specific mesenchyme, but mesenchymal subtypes from pluripotent stem cell sources [77].

The atlas of chromatin accessibility generated here not only provides deeper understanding of fundamental mechanisms underlying genetic control of developmental programs, but also holds relevance to the translational goals of beta cell regeneration and cell replacement therapy. For instance, our comprehensive characterization of chromatin state across endocrine development provides insights into the lineage plasticity observed among endocrine cells [7], and future work can leverage information about active endocrine cell type-specific GRNs to improve strategies for trans-differentiation of non-beta endocrine cells to the beta cell fate. Furthermore, the generation of functionally mature beta cells from hPSCs remains a strong focus of cell replacement therapeutic strategies for patients with diabetes, and such *in vitro* protocols would benefit from an improved understanding of the dynamics in chromatin accessibility across endocrine development *in vivo*. Our dataset identifies which GRNs should be modulated *in vitro* to better approximate *in vivo* development. For instance, it will be interesting to benchmark recently published multi-omic datasets of hPSCs undergoing differentiation to a beta cell fate [78, 79] against our multi-omic dataset generated here to evaluate the fidelity of cells generated *in vitro* to their *in vivo* counterparts.

## ACKNOWLEDGMENTS

The authors would like to acknowledge expert technical assistance from the UCSF Parnassus Flow Cytometry Core, the UCSF Broad Imaging Core, the UCSF Institute for Human Genetics Core, and Dr. Shabrina Amirruddin for assistance with flow analysis. We are grateful to Dr. Evan Deneris for the eFev-EYFP mice.

This work was supported by grants to J.B.S. from The Nora Eccles Treadwell Foundation and the NIH (R01DK118421). S.M.D. was supported by the Kraft Family Fellowship to the UCSF Diabetes Center, the UCSF Discovery Fellows Program, NIH NIGMS IMSD Grant #R25GM056847-23, and NIH/NIDDK diversity supplement R01DK118421-02S1. Z.L. was supported by the Jeffrey G. Klein Family Diabetes Fellowship to the UCSF Diabetes Center.

---

**Supplementary Figure S1:**
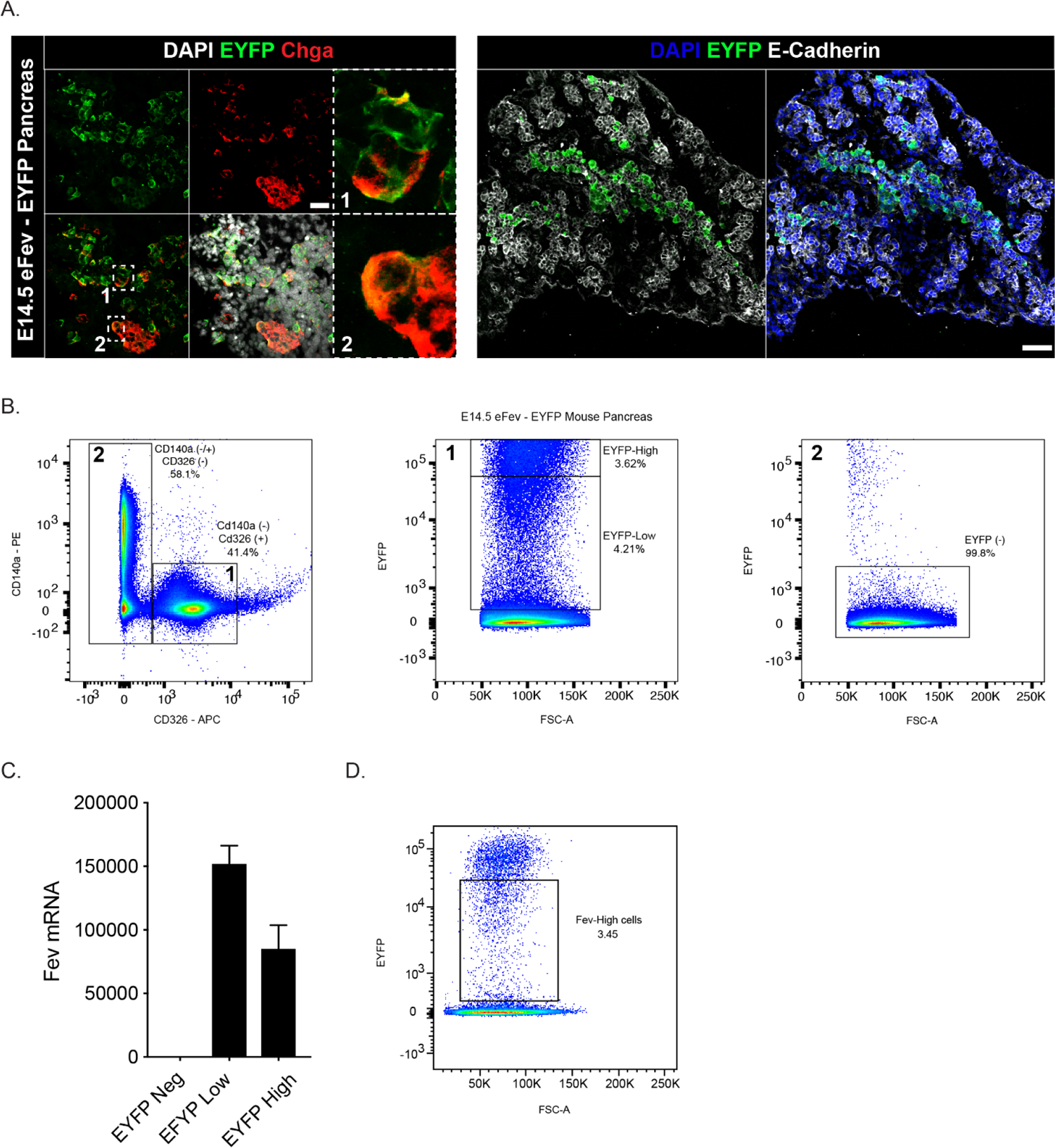
Strategy for enrichment of Fev-expressing pancreatic endocrine progenitor cells. (**A**) Immunofluorescence staining of E14.5 pancreas from eFev-EYFP transgenic mouse embryos. In the left panel, Chga marks differentiated endocrine cells in red, and green reflects EYFP expression. Nuclei are stained with DAPI in white. In the right panel, E-cadherin marks the cell membranes of pancreatic epithelial cells in white, and green reflects EYFP expression. Nuclei are stained with DAPI in blue. (**B**) FACS plots depicting the gating strategy used to assess EYFP expression in eFev-EYFP+ embryos. Cells were dissociated from E14.5 pancreata and stained with antibodies against EpCAM (an epithelial marker) and CD140a (a mesenchymal marker). Two populations were sorted: EpCAM+/CD140-epithelial cells (gate 1) and EpCAM-/CD140+ mesenchymal cells (gate 2). Flow analysis of EYFP expression revealed both an EYFP high and an EYFP low population within the EpCAM+/CD140a-epithelial population, and an absence of EYFP expression in EpCAM-/CD140a+ mesenchymal cells (**C**) Quantitative RT-PCR (Taqman) analysis of sorted EYFP-, EYFP-low, and EYFP-high cells confirmed that EYFP efficiently reflected *Fev* expression in the embryonic pancreas, with no *Fev* expression detected in EYFP-cells. Somewhat higher expression of *Fev* was detected in the EYFP-low vs. -high population. (**D**) FACS plot depicting the gating strategy used for isolating Fev-High (EYFP-low) cells for single nucleus ATAC-Seq (snATAC-Seq). Scale bars in **A** are 20uM (left panels) and 60uM (right panels).

**Supplementary Figure S2:**
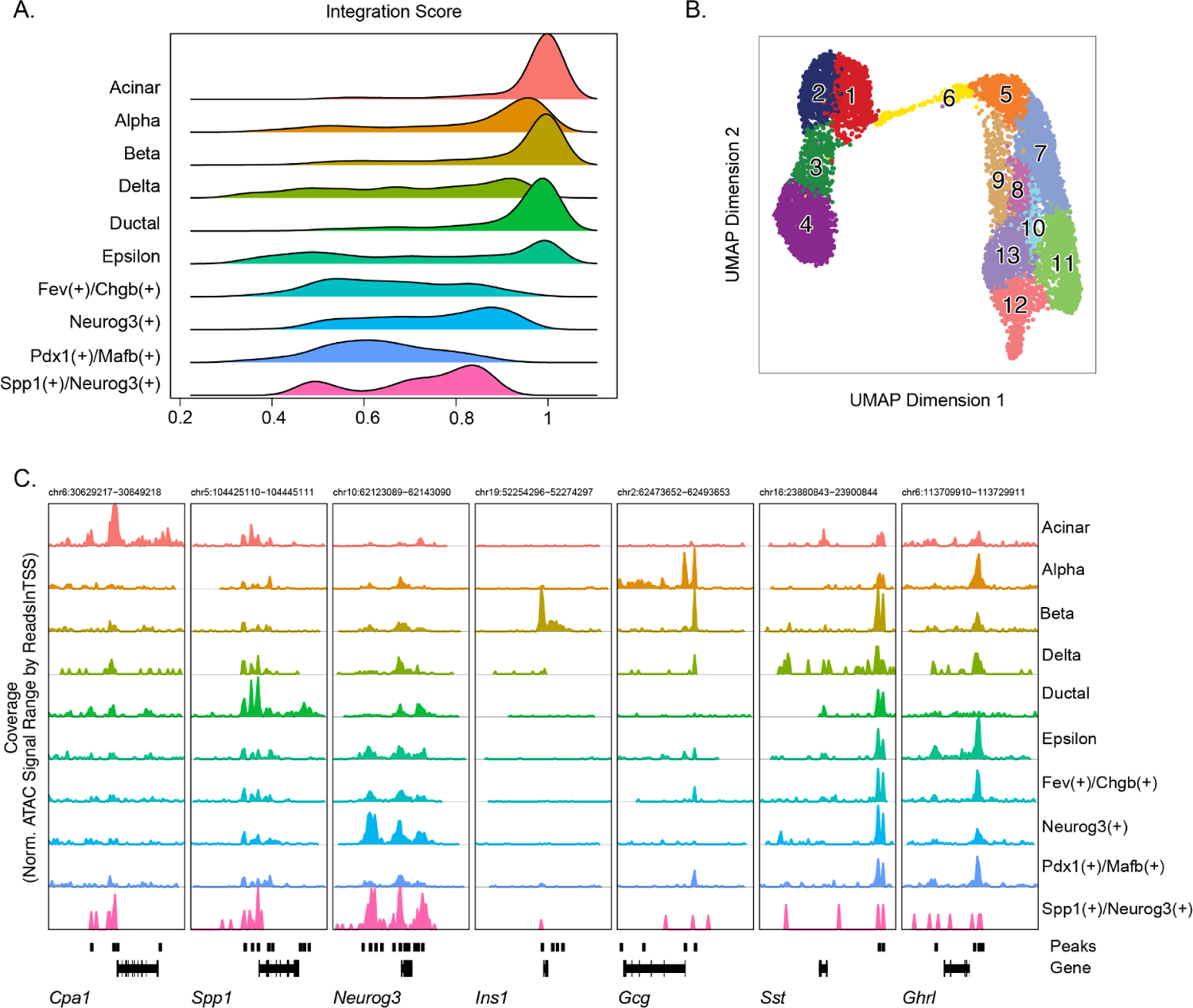
Confirmation of cell type assignments. (**A**) Ridge plots depicting scores from the scRNA-Seq and snATAC-Seq integration for each cell type identified in Figure 2A. A higher integration score indicates higher confidence in the assignment of cell identity. (**B**) UMAP visualization of snATAC-Seq clustering of the epithelial compartment, here without integration with scRNA-Seq data. All clusters in Figure 2A are present even when clustered based on chromatin accessibility alone. (**C**) Track plots of cell type-specific marker genes. Each row indicates a cell type, and each column is a marker gene. The x-axis represents position along the chromosome, which is labeled at the top of each plot. The y-axis represents normalized ATAC signal aggregated across all cells within a population. Regions identified as peaks are depicted as bars above the gene body.

**Supplementary Figure S4:**
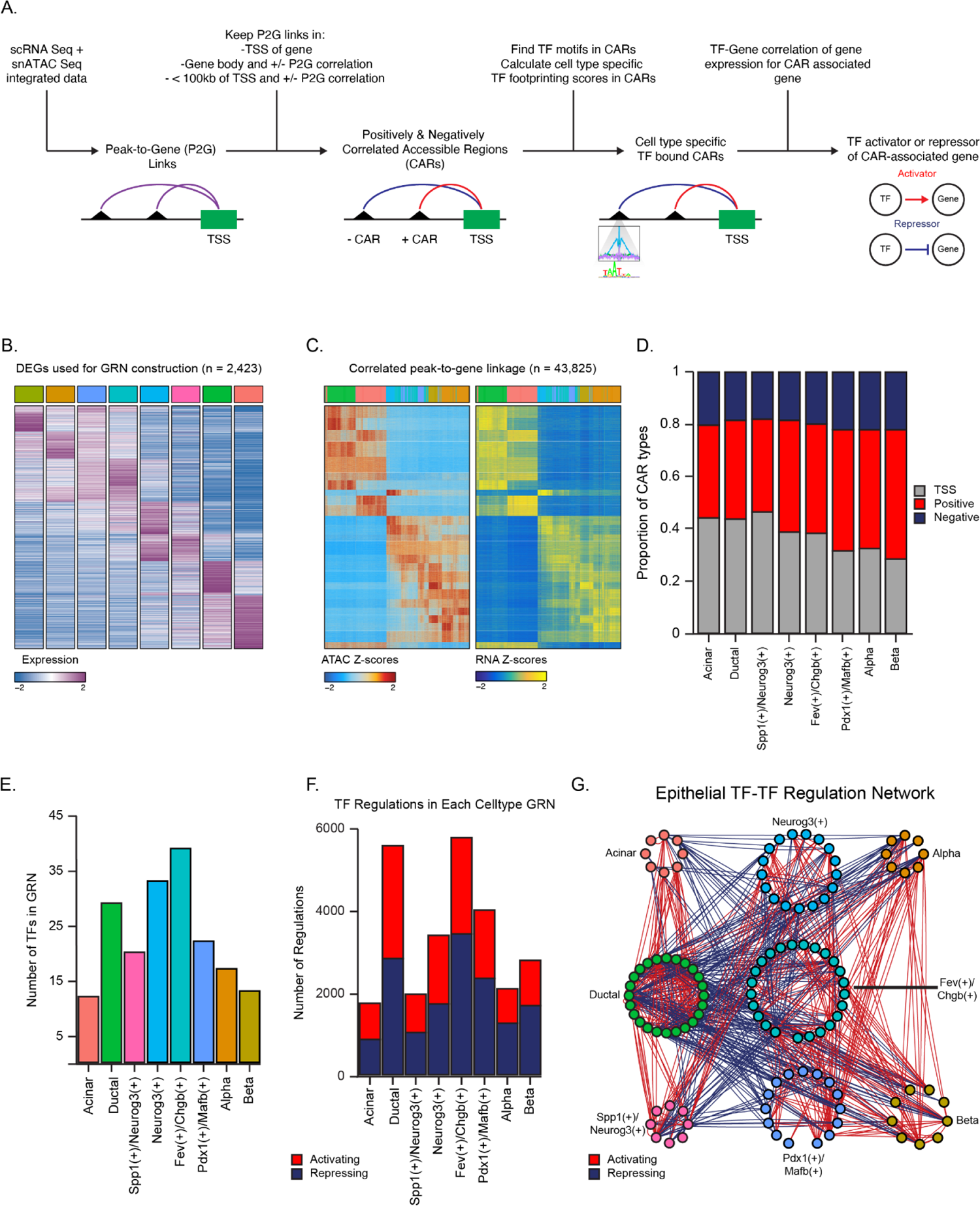
Gene regulatory network map of the developing pancreatic epithelium. (**A**) Diagram depicting the IReNA v2 pipeline. (**B**) Heatmap showing the top differentially expressed genes (DEGs) used as input for GRN construction. Each column represents the average expression of a given DEG within each of the epithelial cell populations shown in Figure 4B. (**C**) Heatmaps showing ATAC accessibility (left) and RNA expression (right) of identified peak-to-gene (P2G) links used as input for GRN construction. Columns represent single cells and rows represent accessible peaks (left) and their correlated genes (right). (**D**) Bar graph cataloging correlated accessible regions (CARs), broken down according to CAR type, for each cell population. The three CAR types include those that lie within the transcription start site (TSS) of a gene (gray), as well as those that are positively (red) or negatively (blue) correlated with their linked gene. (**E**) Bar graph representing the proportion of all TF-gene interactions that are activating vs. repressing. (**F**) Histogram showing the total number of GRN TFs identified within each cell population. (**G**) Bidirectional network diagram depicting all TF-TF interactions between all cell types shown in Figure 4B. TFs are denoted by each node, which is colored by the cell type in which the TF is active in the GRN. Activating regulations are depicted by red lines, while repressing are depicted by blue lines.

**Supplementary Figure S5:**
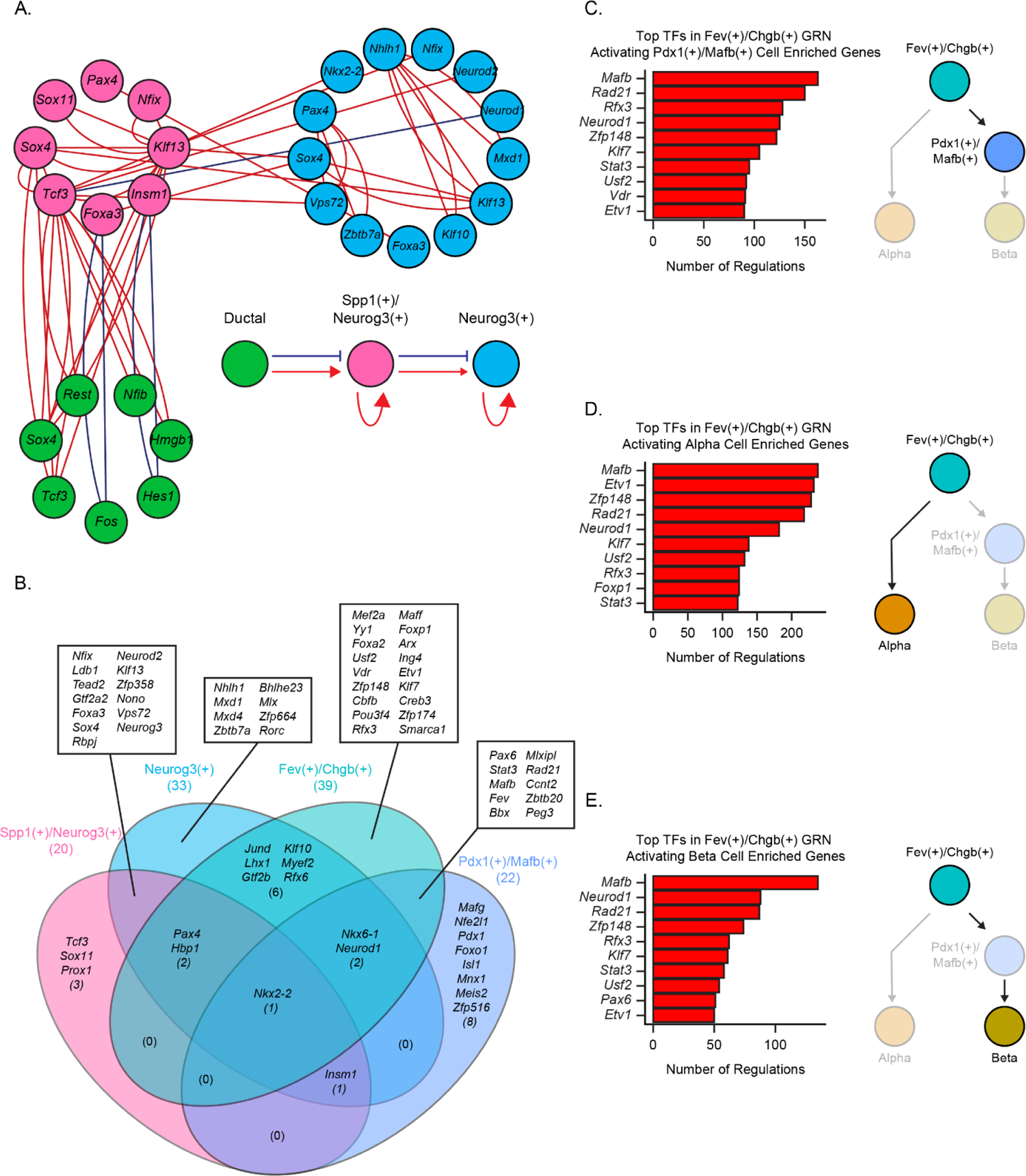
Gene regulatory networks in the endocrine progenitor populations. (**A**) Network diagram depicting the TFs activated (red lines) or repressed (blue lines) among Ductal, Spp1/Neurog3+ and Neurog3+ cell types. Nodes represent TFs and are grouped and colored according to their cell type. The directionality of the regulations are modeled below the diagram. (**B**) Venn diagram depicting overlaps and exclusivity of TFs within the GRN constructed from all four EP populations. (**C-E**) Bar graph depicting the top 10 TFs transcription factors within the Fev/Chgb+ GRN with the highest number of activating regulations of target genes enriched in the Pdx1/Mafb+ (**C**) Alpha (**D**) or Beta (**E**) cell types.

**Supplemental figure 6:**
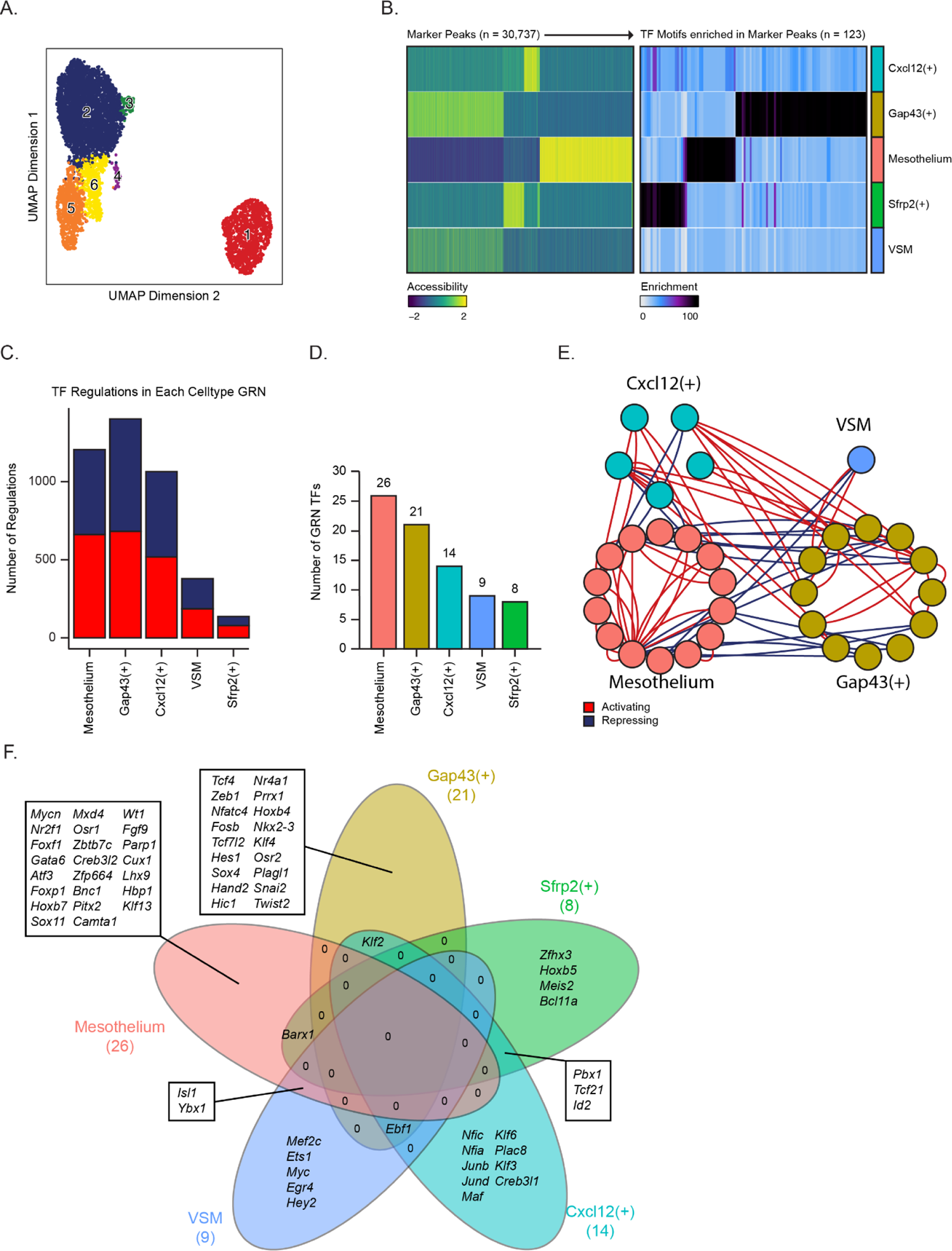
Gene regulatory network in the developing pancreatic mesenchyme. (**A**) UMAP visualization of snATAC-Seq clustering of mesenchymal cells, here without integration with scRNA-Seq data. All clusters in Figure 6A are present even when clustered based on chromatin accessibility alone. (**B**) Heatmaps depict enriched marker peaks (left) and transcription factor (TF) motifs enriched in marker peaks (right) for each mesenchymal cell type. (**C**) Bar graph depicting the proportion of all TF-gene interactions that are activating (red) vs. repressing (blue). (**D**) Histogram showing the total number of GRN TFs identified within each cell population. (**E**) Network diagram showing all TF-TF interactions between all cell types shown in Figure 6A. Each TF is denoted by a node, which is colored by the cell type in which the TF is active. Activating regulations are depicted by red lines, while repressing are depicted by blue lines. (**F**) Venn diagram depicting shared and exclusive Mesenchymal TFs within the GRN. VSM, vascular smooth muscle.

